# OmniScore: A geometric foundation encoder for universal biomolecular representation and scoring

**DOI:** 10.64898/2026.08.28.747942

**Authors:** Tien-Cuong Bui, Yiyu Hong, Jinung Song, Yunwen Tao, Injae Chung, Jun Gyou Park, Junsu Ko, Juyong Lee

## Abstract

Biomolecular structures provide a common physical basis for understanding molecular function and interactions. However, computational models typically specialize in individual molecular classes or tasks, hindering a unified representation of structural information across biological systems. Here we introduce OmniScore, a universal geometric encoder that learns shared representations of proteins, nucleic acids, small molecules and their complexes. Pretraining integrates geometric graph encoding, sequential modeling and tokenization with structural and physicochemical supervision. The frozen pretrained encoder supports complex-level model quality assessment and interaction prediction, as well as residue-level prediction, through task-specific heads. OmniScore outperforms the evaluated antibody–antigen quality-assessment baselines and also improves RNA structure ranking over baselines. For peptide–major histocompatibility complex interaction prediction, the OmniScore embeddings complement structure-prediction confidence metrics to improve predictive performance. Together, these findings support OmniScore as a shared geometric foundation for biomolecular representation and scoring across molecular classes and prediction scales, which may facilitate multi-component therapeutics development.

## 1 Introduction

Biomolecular complexes connect structure to function. Proteins, nucleic acids, small molecules, antibodies, and their combinations form the physical interfaces that control recognition, catalysis, signaling, and therapeutic response. Scoring these complexes is therefore a central problem in computational biology and drug discovery: given a three-dimensional structure or a set of candidate structures, we need to estimate whether the interaction is plausible, how strong it is, and which model should be trusted. These assessments, together with residue-level prediction, depend on representations that capture local chemical geometry and broader structural context. However, learning such representations across molecular classes remains challenging. Molecular systems differ in size, chemistry, flexibility, and available supervision, while experimental complex structures and affinity labels remain unevenly distributed across interaction types. As structure prediction models continue to generate increasingly diverse complexes, the need for reusable structural representations that support scoring and other prediction tasks across molecular classes becomes more urgent.

Despite significant progress, most methods still solve the problem in isolation. Protein–ligand scoring functions such as AK-Score [1, 2] and GenScore [3] focus on binding affinity, pose ranking, docking, or screening tasks in protein–ligand systems (e.g., Comparative Assessment of Scoring Functions 2016, CASF-2016 [4]). Protein–protein methods such as GNN-DOVE [5] and ProAffinityGNN [6] evaluate docking decoys or predicted complex quality, while antibody–antigen methods target pose identification [7], epitope prediction [8, 9], and antibody–antigen DockQ estimation [7, 10]. Lately, more general biomolecular interaction models, such as Generalist Equivariant Transformer (GET) [11] and BioScore [12], have moved toward unified geometric representations across molecular domains. These directions are promising, yet a practical gap remains: system-specific models often encode each molecular class with task-tailored assumptions, while general scoring approaches still need representations that preserve both local chemical geometry and transferable structural context across proteins, nucleic acids, and small molecules. Using only a task-specific supervised head can also make learning heavily dependent on sparse labels.

We introduce OmniScore, a pretrained geometric encoder for modeling and scoring the structures of diverse biomolecular complexes. Our core idea is to learn a common latent space whose pretrained representations can be reused for downstream prediction while keeping the encoder frozen. To achieve this goal, we train the encoder within an autoencoder framework, with a tokenizer and a reconstruction decoder as components of pretraining. Specifically, the encoder learns geometry-aware representations of different complexes by first converting each input structure into two coupled views: a two-level heterogeneous graph and an aligned list of atoms. By combining an equivariant graph transformer (EGT) with a bidirectional Mamba, the encoder efficiently merges geometric and long-range sequential features. Additionally, it incorporates a quantizer as a feature bottleneck module, based on finite scalar quantization (FSQ) with residual coding [13, 14], to encourage compact representations of common patterns or interactions. The decoder is specifically for reconstructing 3D coordinates given quantized latent embeddings, crucial for the pretraining process. Downstream heads can access graph, continuous sequence, or quantized representations, making the pretrained encoder the common basis for different prediction tasks.

We pretrained OmniScore with multiple objective functions, including structure reconstruction, token corruption recovery [15, 16], and various other auxiliary objectives related to property prediction. To achieve the goal, we collected and curated multiple datasets, including monomers, complexes of proteins, nucleic acids, and small molecules. After pretraining, we froze the encoder and quantizer and trained only task-specific prediction heads for downstream tasks, including DockQ prediction for antibody–antigen structures and binding affinity prediction for protein–ligand complexes. To assess the utility of these frozen representations at both the residue and structure levels, we conducted experiments on the residue-level evaluations of StructTokenBench [17], antibody–antigen structure quality assessment [7], RNA structure ranking (RSR), peptide–major histocompatibility complex (pMHC) interaction classification, and CASF-2016 protein–ligand scoring and ranking benchmarks [18]. Furthermore, we also performed rigorous ablation studies of various model configurations to understand the effect of added features and components to find the most stable configurations. Across these evaluations, frozen OmniScore representations supported competitive functional-site prediction and antibody–antigen quality assessment, while a GNN head on graph representations improved RNA structure ranking over retrained ATOM3D baselines. In pMHC classification, graph representations complemented structure-prediction confidence features to improve predictive performance. The results further indicate that downstream utility depends on the representation stage and that better reconstruction does not necessarily translate into better downstream performance.

## 2 OmniScore Geometric Encoder

### 2.1 Datasets

We collected datasets categorized into two groups: complex datasets and monomer and small molecule datasets. Complex datasets provide cross-chain and cross-molecule interactions, while monomer and small molecule datasets provide atomistic packing examples allowing the model to learn atomic-level interactions.

#### Complex Datasets

For complex datasets, we include PDBbind v2020 [18] and the antibody/nanobody–antigen dataset used in IgPose [7]. PDBbind v2020 includes curated protein–protein, protein–ligand, nucleotide–ligand, and protein– nucleotide complexes. These complexes provide binding affinity labels, including the dissociation constant (*K*_*d*_), the inhibition constant (*K*_*i*_), and half-maximal inhibitory concentration (IC_50_) when available. We use *pK* as a collective notation for the negative base-10 logarithm of the available affinity measure expressed in molar units. The IgPose dataset includes the crystal structures of antibody/nanobody–antigen complexes and their predicted structures through Chai-1 [19] and Boltz-2 [20]. Note that not all crystal structures have the predicted structures due to computation errors (e.g. out of memory).

#### Monomers and Small Molecules

These datasets include CATH, CASP14, CASP15, RNA3DB, and NablaDFT, adopted from Bio2Token’s code repository [21]. We used only the processed local splits available in the code repository. We also created monomer and small molecule versions of the complex dataset by extracting individual chains and ligands from PDB files and denoting them PDBbind Protein Monomers, PDBbind Small Molecules, and IgPose Crystal Monomers. These subsets provide additional proteins, nucleotides and small molecules for the reconstruction pretraining process.

#### Data Split Policies

We use different split policies according to the source dataset. PDBbind v2020 complexes were randomly split into training and validation sets with a 7:3 ratio and used only in the pretraining phase. The PDBbind-derived monomer datasets inherit this split from their source complexes. Thus, protein chains, ligands, and nucleotide chains from a validation complex remain in the validation partition. The structures from the IgPose dataset inherit the original antigen-clustered split policy. IgPose groups complexes by antigen-clustered identifiers to prevent antigen-level leakage across train, validation, and test partitions. CATH, RNA3DB, and NablaDFT use the processed splits provided by the Bio2Token repository [21]. As additional structure data, submitted model structures from CASP14 and CASP15 were also added to the training partition during the pretraining phase. For both SID-R and CASF-2016, we used protein sequence identity to remove redundancy between the benchmark and training datasets.

#### Pretraining Data Statistics

The statistical information of datasets used in the pretraining phase is listed in Table 1. The effect of excluding ligand-containing structures from pretraining is evaluated in Supplementary Fig. S1. OmniScore is pretrained with a mixture of complexes and monomers. As we aim to construct a common latent space of diverse molecular structures, different input modalities provide complementary structural contexts. For instance, complexes provide long-range and cross-entity interactions, while monomers provide isolated local geometry without interface information. Additionally, small molecules can help the deep learning model learn covalent bond and angle patterns, which are required for ligand coordinate reconstruction. Consequently, no test complex or close homologue contributed structures during pretraining.

**Table 1:**
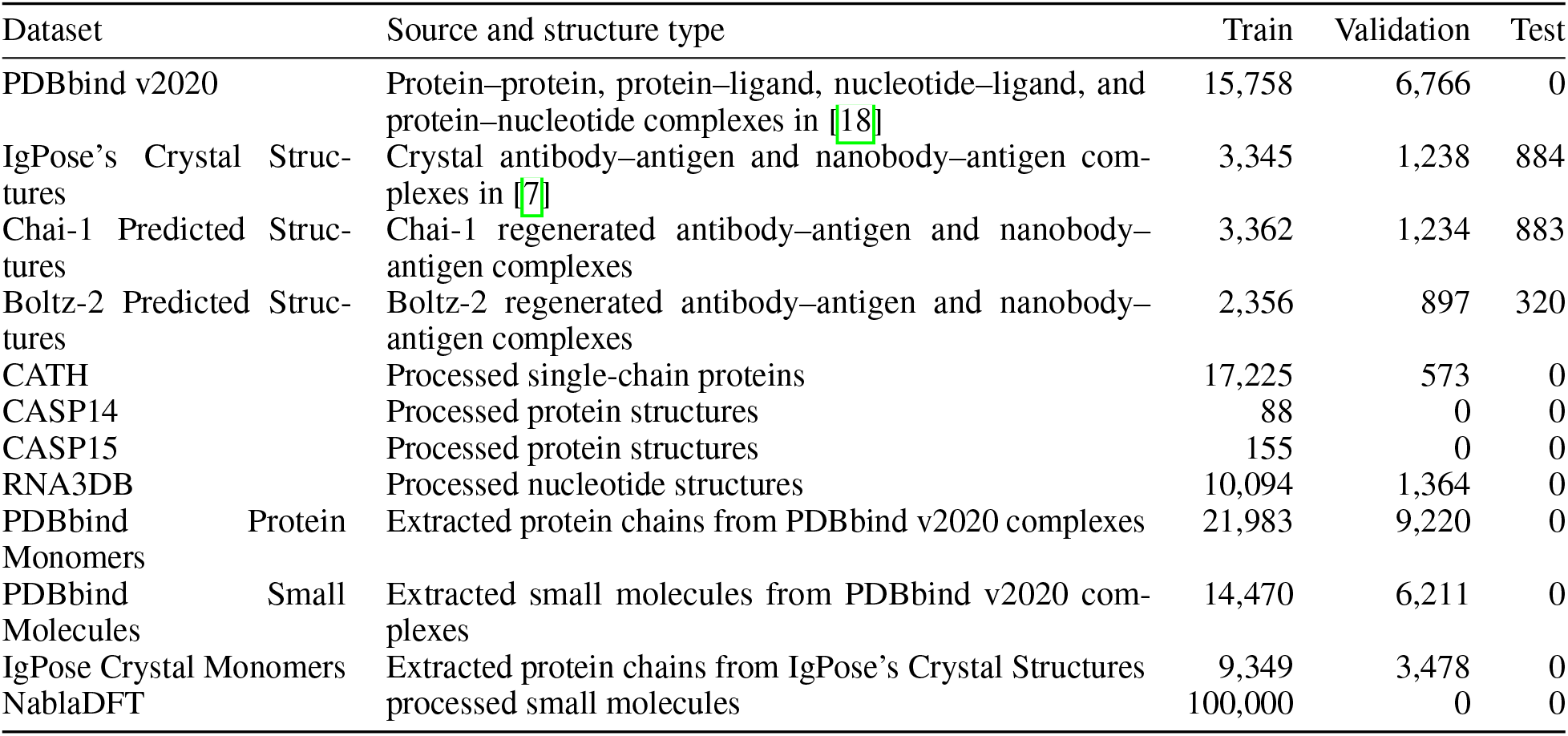
Datasets used in OmniScore pretraining and finetuning.

**Table 2:** Datasets and partitions used for downstream evaluation. DockQ prediction uses the antigen-clustered SID-R split adopted from IgPose. References identify the source datasets and benchmark protocols.

| Task | Training and validation | Benchmark | Target |
| --- | --- | --- | --- |
| DockQ prediction | IgPose crystal and Chai-1/Boltz-2 predicted structures [7] | SID-R test split | DockQ |
| Binding affinity scoring | PDBbind v2020 [18] | CASF-2016 [4] | $pK$ |
| RNA structure ranking | ATOM3D RSR [22]: RNA-Puzzles candidates after RNA3DB overlap removal | RNA Puzzles 14 (bound/free), 18, and 21 | All-atom RMSD |
| pMHC interaction classification | PREpiBind Qualitative [25]: 90,328 training and 22,543 validation complexes (Fold 0) | PREpiBind Qualitative test partition (48,352 complexes; 30%) | Binding/non-binding |
| StructTokenBench | StructTokenBench datasets [17] residue-level | Benchmark splits | Residue-level labels |

#### Finetuning for downstream tasks

We evaluate five downstream settings: antibody/nanobody–antigen DockQ prediction, RNA structure ranking, pMHC interaction classification, protein–ligand binding affinity scoring, and StructTokenBench residue-level prediction. For DockQ prediction, we follow the IgPose problem setting, where native structures are assigned DockQ scores of 1, and regenerated structures provide the continuous DockQ scores ranging over [0, 1]. For protein–ligand binding affinity prediction, we use PDBbind v2020 as the training/validation sets and CASF-2016 as the benchmark set. We adopt StructTokenBench residue-level benchmark tasks to evaluate physical and functional information contained in the frozen OmniScore residue embeddings.

For RNA structure ranking (RSR), we use the released ATOM3D candidates [22]. Each example is a FARFAR2-generated candidate [23] for an RNA-Puzzles target [24], labelled by its all-atom root-mean-square deviation (RMSD) to the experimental structure. We checked for overlap between the RNA3DB pretraining dataset and RNA-Puzzles data and removed shared structures before the RSR evaluation. The evaluated RNA targets do not overlap with data used to train either the pretrained encoder or the prediction heads. The evaluation comprises RNA Puzzles 14, 18, and 21, with bound and free forms of Puzzle 14 treated as separate structure groups. All chains are retained for multi-chain targets.

For pMHC interaction classification, we use the Qualitative dataset provided in the PREpiBind study [25], which was curated from the Immune Epitope Database (IEDB; accessed 20 April 2025). The source dataset comprises 161,223 peptide–MHC class II entries (58.1% positive), restricted to linear, unmutated 15-mer peptides paired with human HLA-DR, HLA-DQ, or HLA-DP, or murine H2 molecules. Labels indicate experimentally observed binding or non-binding, and performance is evaluated by the area under the receiver operating characteristic curve (AUROC). We reuse the original PREpiBind data partitions: a 70:30 training–test split stratified by class label and MHC serotype, with

Fold 0 of the training partition used for validation. The resulting sets contain 90,328 training, 22,543 validation, and 48,352 test complexes (approximately 58% positive). Entries sharing an identical peptide and MHC *β*-chain sequence are grouped in the same training or test partition to prevent overlap of identical binding contexts.

### 2.2 Input Preparation

Vectors, matrices, and feature tensors are shown in bold; scalars, indices, sets, and functions are unbolded. As OmniScore supports diverse biological modalities, such as proteins, RNA, and small molecules, we abstract molecular structures into blocks belonging to one of the following types: ℬ = {residue, nucleotide, atom}. Each residue or nucleotide block can contain multiple atom rows, whereas an atomic block represents one heavy atom from a small molecule. This block representation preserves biological units for macromolecules while retaining atom-level granularity for small-molecule chemistry. Essential architecture notation is summarized in Supplementary Table S1, and block layouts and edge construction are detailed in Supplementary Section S2.

Based on the block abstraction, OmniScore builds a heterogeneous graph *G* and a flattened atom-wise sequence *S* for an input structure. The graph provides sparse spatial neighborhoods at block level, while the sequence preserves atom-wise order and atom/block identities for sequential encoding. Specifically, *G* = (*V, ε,T, R*, **X**), where is a set of blocks, is a set of typed block-to-block edges, *T* contains atom and block token attributes, *R* contains block-type and edge-type attributes, and **X** contains atomic coordinates. We define the ordered atom-wise sequence 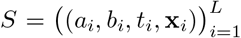, where *a*_*i*_ is an atom token, *b*_*i*_ is a block token, *t*_*i*_ *∈ B* is the block type, and **x**_*i*_ *∈* ℝ^3^ is the coordinate of atom *i*. The atomic block type *t*_*i*_ should not be confused with an atom token *a*_*i*_, where an atomic block defines graph granularity, while an atom token defines chemical identity such as carbon, nitrogen, or oxygen.

### 2.3 Pretraining OmniScore

#### Overall Pipeline

We design the pretraining pipeline as an autoencoder including an equivariant graph transformer and a bidirectional Mamba encoder, followed by a token quantizer and a Mamba decoder (Fig. 1). The overall pipeline can be briefly written as follows: 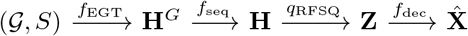. Specifically, the encoder, defined as *f*_enc_(*G, S*) = (*f*_seq_ ° *f*_EGT_) (*G, S*), performs two-phase encoding via an equivariant graph transformer (EGT) and a bidirectional Mamba encoder to capture both geometric and sequential information. The encoded features **H** (pre-quantization embeddings) are then passed to a residual finite scalar quantization (RFSQ) module *q*_RFSQ_ to output **Z**–quantized latent embeddings. Finally, a bidirectional Mamba decoder *f*_dec_ reconstructs atom coordinates 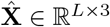 from **Z**. The decoded coordinates 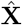 are supervised by geometric reconstruction losses against **X** = [**x**_1_, …, **x**_*L*_]^*⊤*^. Quantized latent representations **Z** are also used by auxiliary objective functions and downstream finetuning procedures. The tokenizer and decoder support encoder pretraining, while downstream heads use graph representations **H**^*G*^, continuous sequence embeddings **H**, or quantized embeddings **Z** according to the task. Equivariance refers to the graph module; the full encoding pathway is described as geometry-aware.

**Figure 1:**
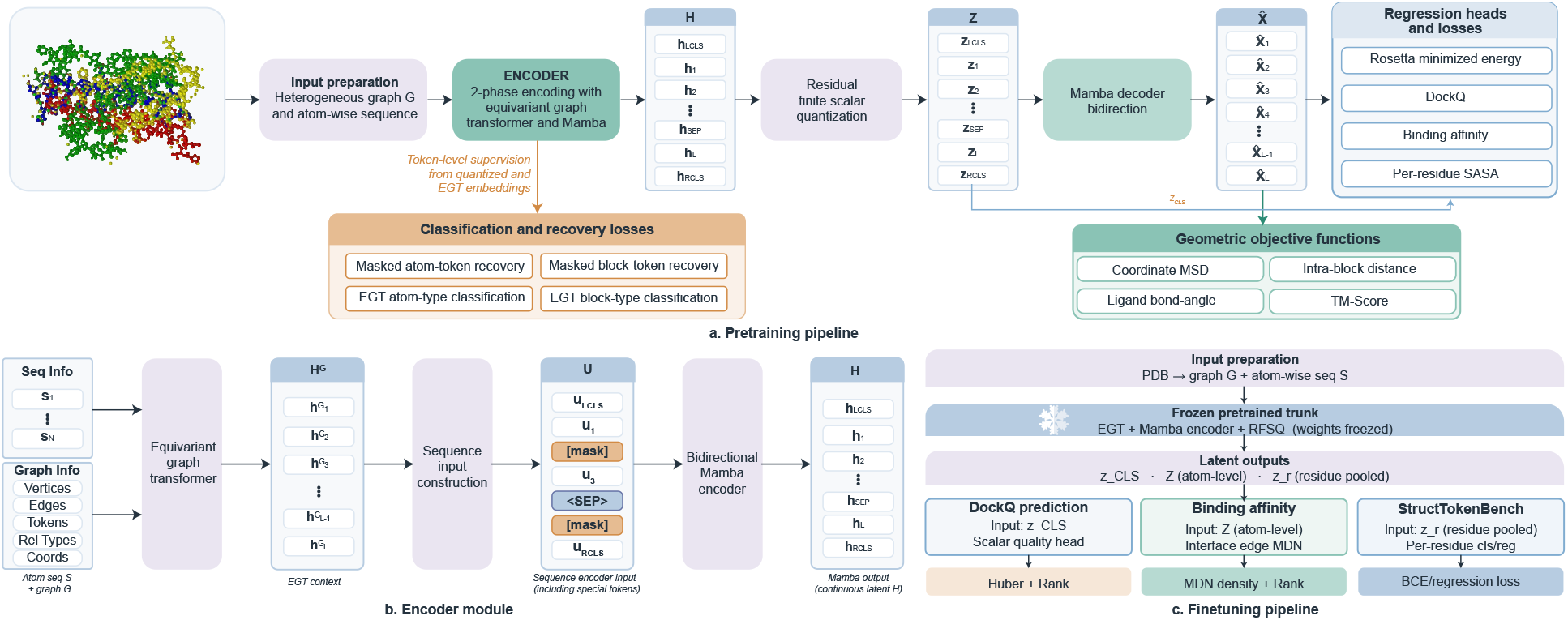
Overview of the OmniScore geometric encoder and training workflow. (A) Pretraining pipeline. Each molecular structure is converted into a heterogeneous graph *G* and an atom-wise sequence *S*, encoded by a two-phase encoder combining an equivariant graph transformer (EGT) and bidirectional Mamba, quantized by residual finite scalar quantization (RFSQ), and decoded by a Mamba decoder to reconstruct coordinates 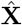. Orange boxes denote auxiliary token-level objectives, and green boxes denote geometric reconstruction objectives. Structure-level regression heads use the global classification (CLS) representation **z**_CLS_ for Rosetta minimized energy, DockQ, and binding-affinity prediction. (B) Encoder module. Sequence information and graph information are first processed by EGT to produce aligned geometric context **H**^*G*^ then mapped back to atom-wise in the sequential order; the input token sequence **U**, including masked positions and special separator/classification tokens, is then passed through a bidirectional Mamba encoder to produce continuous embeddings **H**. (C) Finetuning pipeline. The pretrained OmniScore encoder converts Protein Data Bank (PDB) inputs into latent outputs used by task-specific heads for DockQ prediction, binding affinity with an interface mixture-density network (MDN), and StructTokenBench residue-level classification or regression. BCE denotes binary cross-entropy.

#### Equivariant Graph Transformer

The equivariant graph transformer accepts the heterogeneous graph *G* as input and outputs geometry-aware features aligned with the atom-wise sequence *S* for the subsequent sequence encoder. For each block *u ∈ V*, EGT first maps its atom tokens and block token to embedding spaces.

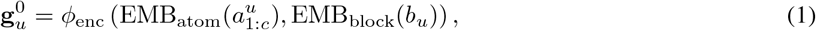

where EMB is the embedding lookup function, *c* is the number of atom channels for block *u*, and *ϕ*_enc_ concatenates each atom-channel embedding with its parent block embedding. For layer *ℓ*, EGT updates atom and block features by

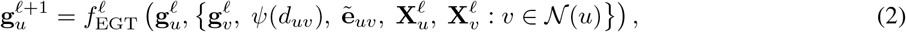

where 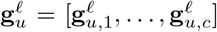 denotes the channel-wise feature tensor of block *u, N* (*u*) = *{v 2 V* : *{u, v} ∈ ε}* is the set of neighboring blocks of *u, ψ* (*d*_*uv*_) is a radial basis encoding of distance, 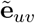 is an edge-type embedding, and 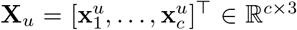 contains the atom-channel coordinates of block *u. f*_EGT_ can be any geometric graph neural network (GNN) satisfying equivariance of the input coordinates. In the current version, we adopt Generalist Equivariant Transformer (GET) [11], the geometric transformer model supporting hierarchical representation of biological structures. Therefore, EGT can operate on normalized biological units while still exposing atom-channel geometry inside each block via the two-level block-atom representation.

#### Sequence Encoder

To support sequential encoding, the EGT output is mapped from block-channel features to atom-wise features. For a graph with *N* blocks and a sequence with *L* atom rows, we extract the corresponding EGT’s atomic feature vectors to construct 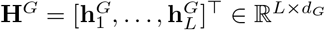. For each atom row *i*, its coordinate **x**_*i*_, atom embedding, EGT feature 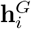, and block embedding are concatenated to form the sequence input as follows:

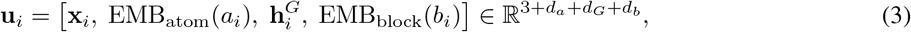

where the EMB_atom_ and EMB_block_ functions with dimensions of *d*_*a*_ and *d*_*b*_ are adopted from EGT (eq. 1) to maintain token-embedding consistency and reduce the number of learned parameters. To handle multi-entity inputs and support structure-level downstream tasks, we introduce entity-separation and classification tokens. A separator (SEP) token ⟨ sep ⟩ separates chains or molecular entities and is masked out during coordinate reconstruction. Left and right classification tokens, ⟨LCLS⟩ and ⟨RCLS⟩, are placed at the beginning and the end of the sequence. The sequence input **U** = [**u**_LCLS_, **u**_1_, …, **u**_SEP_, …, **u**_*L*_, **u**_RCLS_]^*⊤*^ is processed by a bidirectional Mamba encoder. To align these special tokens with the same feature layout as atom rows, their token-embedding dimensions are learnable, while their coordinate and graph-context components are initialized according to their roles. For the two classification tokens, the coordinate component is initialized with the structure centroid obtained by averaging over all **x**_*i*_, and the graph-context component is initialized with the pooled EGT representation obtained by averaging over all 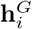. Special tokens such as ⟨ sep ⟩ and the padding token ⟨ PAD ⟩ are excluded from both averages. The remaining token-embedding components are learned parameters in 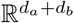. The classification tokens are also passed through the quantizer and later used for structure-level regression heads. These tokens are excluded from structure reconstruction. This separation prevents global property prediction and coordinate reconstruction from competing for the identical output positions.

#### Masking Paradigm

OmniScore employs a selectable denoising paradigm for the pretraining phase. The model uses token masking and optional coordinate masking. Here, we describe a general masking paradigm because these choices are interchangeable training options. Token and coordinate corruption rules are detailed in Supplementary Section S4.5. Let Ω_atom_ denote the set of atoms comprising the input molecular structure, indexed by their positions in the flattened sequence; special tokens and padding are excluded. The mask set ℳ ⊆ Ω_atom_ is sampled independently or using contiguous block spans:

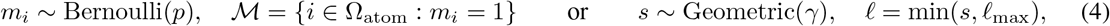

where *p* is the probability of a random masking process [15], *ℓ* is the desired length of a span-based masking algorithm [16], *ℓ*_max_ is the maximum span length, and is 0.2, as in [26].

#### Residual Finite Scalar Quantization

We apply RFSQ to the continuous encoder representations **H**, combining finite scalar quantization (FSQ) [13] with residual coding [14]. Each vector **h**_*i*_ *∈* ℝ^*d*^ is projected to a low-dimensional space, **v**_*i*_ = **W**_in_**h**_*i*_, and quantized in successive stages that encode the remaining residual. The combined codes are projected back to the model dimension as **z**_*i*_ = **W**_out_ RFSQ(**v**_*i*_). We use **z**_CLS_ = **z**_LCLS_ + **z**_RCLS_ for global prediction and **Z** = [**z**_1_, …, **z**_*L*_]^*⊤*^ for residue-wise tasks and structure reconstruction.

#### Decoder

To reconstruct the input structure, the decoder is implemented based on the Mamba architecture. It receives only the quantized atom-wise sequence **Z** and outputs predicted coordinates 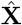 in the same order as the input atomic sequence:

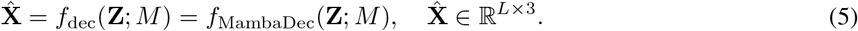

#### Pretraining Objective Functions

The general pretraining objective includes the reconstruction loss term *L*_geo_, the fine-grained classification term *L*_cls_, and the auxiliary loss term *L*_aux_, which can be formalized as follows:

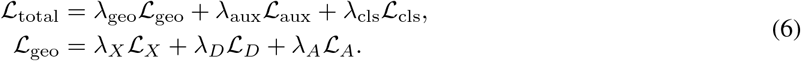

Coordinate reconstruction loss *L*_*X*_ minimizes atom-wise coordinate errors, while the intra-block distance loss *L*_*D*_ constrains local geometry:

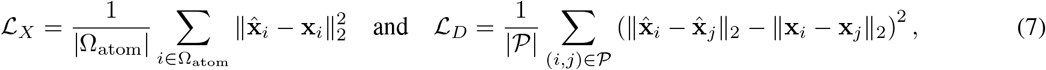

where *P* denotes the set of intra-block atom pairs and Ω_atom_ denotes the set of atoms in the input biomolecular structure. We define the angle loss function over the set *T* of bonded atom triplets (*i, j, k*) derived from bond edges to constrain the geometry of small molecules:

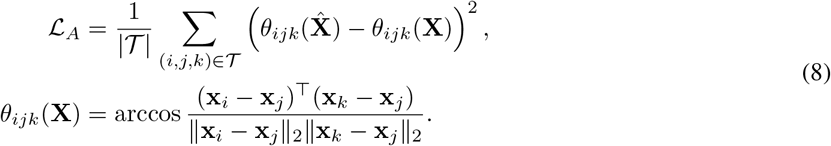

The auxiliary regression loss aggregates the available auxiliary global property prediction targets using a label mask and the Huber loss:

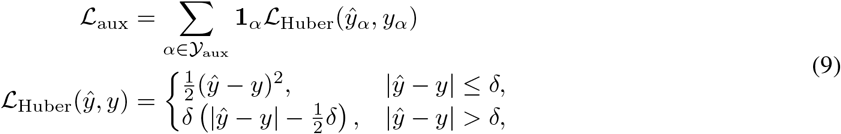

where *Y* _aux_ includes Rosetta minimized energy, DockQ, protein–ligand and protein–protein binding affinity values (*K*_*d*_, *K*_*i*_, and IC_50_), and per-residue solvent exposure magnitudes, including solvent accessible surface area (SASA) and half-sphere exposure (HSE), while **1**_*α*_ masks samples without the auxiliary label *α*.

The fine-grained classification loss combines two token-level supervision paths:

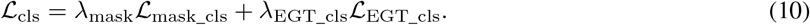

Here *L*_mask_cls_ supervises recovery of corrupted atom and block tokens from masked quantized embeddings, and *L*_EGT_cls_ supervises atom and block type prediction from the EGT output embeddings 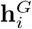. Both terms are cross-entropy losses over the atom-token vocabulary and the block-token vocabulary. Note that *A* weights control the contributions of loss terms, with weights ranging from 0.01 to 0.1 (Supplementary Table S4).

### 2.4 OmniScore model variants

We evaluated seven OmniScore model variants, ranging from approximately 24M to 124M parameters, by varying the hidden dimension and the depths of the Mamba encoder and decoder. The hidden dimension was either 512 or 1024, with encoder depths of 8 or 12 layers and decoder depths of 4 or 6 layers. Model names indicate approximate parameter counts; the hidden dimensions and Mamba encoder/decoder depths for each variant are summarized in Supplementary Table S2. Parameter counts by component are provided in Supplementary Table S3.

### 2.5 Finetuning OmniScore for downstream tasks

For DockQ prediction, RNA structure ranking, pMHC classification, protein–ligand scoring, and StructTokenBench evaluation, the pretrained encoder and quantizer remain frozen and only the task-specific prediction modules are optimized. This setting keeps the structure representation fixed and limits the number of trainable parameters for downstream datasets. Without further explanation, we configure a 2-layer multilayer perceptron (MLP) having the same hidden dimension size as the encoder as a default regression network for DockQ prediction and binding affinity prediction tasks. Software, optimization settings, and implementation details are provided in Supplementary Section S4.

#### DockQ Prediction

The DockQ head predicts the quality of predicted antibody–antigen and nanobody–antigen structures. The finetuning objective combines the Huber regression and correlation-based ranking losses:

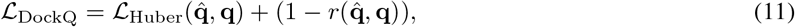

where 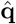 and **q** collect predicted and true DockQ values over a minibatch, *L* _Huber_ averages the element-wise Huber losses, and *r* denotes the Pearson correlation function.

#### RNA Structure Ranking

We use the pretrained OmniScore parameter version v3 and keep the backbone parameters frozen for all reported RSR experiments. Nucleotides are represented by nine-atom blocks processed by the EGT [11], followed by the Mamba encoder [27] and scalar quantization [13], as described above. We compare MLP and ridge heads on output embeddings, linear probes on intermediate representations, and a residue-graph GNN.

For each structure group, we compute the Spearman rank correlation (RS), *ρ*, between the method score (predicted RMSD for learned regressors) and true RMSD over its candidate structures. The primary metric gives each of the three RNA targets equal weight by first averaging the bound/free correlations for Puzzle 14:

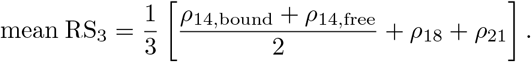

We also report the four-group mean, mean RS_4_ = (*ρ*_14,bound_ + *ρ*_14,free_ + *ρ*_18_ + *ρ*_21_)*/*4, which gives the two Puzzle 14 groups separate weights. Both metrics average within-group correlations; neither is a pooled correlation across candidates from different RNAs. Larger values indicate better ranking of candidates for the same RNA. Representation, probe variant, ridge strength, training epoch, and objective are selected by validation mean RS; test data are used only for evaluation.

The post-quantization MLP takes a 1,024-dimensional concatenation of post-quantization CLS and mean-pooled post-quantization embeddings and has layer widths 1024 → 256 → 256 → 1. It is trained on standardized RMSD with 0.3 *L*_MSE_ + 0.7(1 *r*), where *L*_MSE_ is mean squared error and *r* is Pearson’s correlation within a minibatch of 64 candidates from one target. Predictions from five seeds are ensembled. Additional MLP variants use either mean-pooled pre-quantization embeddings or pre-quantization CLS embeddings. We also evaluate ridge regression on the concatenated post-quantization CLS and mean embeddings.

Linear probes fit ridge regression to pooled EGT and Mamba representations, with regularization strength *α ∈ {*10^*-*1^, 10^0^, …, 10^5^*}*. The centered variant removes each training target’s feature and RMSD means before fitting:

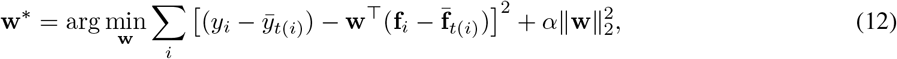

where **f**_*i*_ is the pooled feature vector, *y*_*i*_ is the true RMSD, *t*(*i*) identifies its target, and bars denote training-target means. Candidates are scored by (**w**^*\**^)^*⊤*^ **f**_*i*_. The global variant fits standardized RMSD without target centering. Validation selects the representation, centering variant, and regularization strength.

For the residue-graph GNN, each nucleotide node receives a 128-dimensional EGT feature averaged over its nine-atom block. Edges connect the 10 nearest block centroids and carry radial-basis distance and sequence-separation features. Three message-passing layers with 128 hidden units are followed by mean/max pooling and an MLP. Predictions from three seeds are ensembled. Validation selects the within-target objective over the global objective for the pretrained checkpoint:

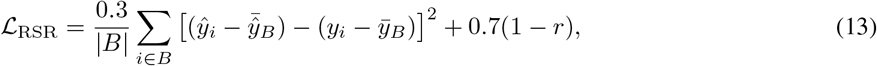

where *B* is a minibatch containing candidates from one target, *ŷ*_*i*_ is its predicted RMSD, and 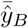 and 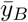 are batch means. Here, *r* is Pearson’s correlation between predictions and targets within *B*.

We evaluate the FARFAR2 Rosetta total score [23] and the individual Rosetta terms `fa_stack, N_BS`, and `hbond_sc`. ARES (Atomic Rotationally Equivariant Scorer) is evaluated with its released weights [28]. The ATOM3D three-dimensional convolutional neural network (3DCNN) and GNN baselines [22] are retrained on RSR and evaluated as three-seed ensembles on the same four evaluation groups. Supplementary Table S5 reports three-target, four-group, and per-group correlations.

#### pMHC Interaction Classification

Each pMHC complex comprises the MHC-II *α* and *β* chains and a 15-mer peptide. We generated one structure per complex with Boltz-2, using MSAs for the MHC-II chains, no peptide MSA, three recycling steps, and 200 diffusion steps. The pretrained OmniScore-24M encoder remained frozen and embedded each complex once. We compared three representation stages: 128-dimensional EGT outputs **H**^*G*^ before Mamba (G), 512-dimensional continuous Mamba outputs **H** before quantization (P), and 512-dimensional post-quantization embeddings **Z** (E). An 8 Å peptide–MHC contact cutoff defined the interface. MLP classifiers used mean-pooled interface atom embeddings, whereas graph attention network (GAT) classifiers used residue-averaged embeddings on a *k* = 10 nearest-neighbour graph of interface residue centroids. We trained only the downstream classifiers, with and without 16 Boltz-2 confidence features, and evaluated AUROC on the full test set. Model selection used validation AUROC; all runs used seed 0. Baselines comprised a confidence-only classifier and same-test-set results for NetMHCIIpan-4.3 and MixMHC2pred-2.0. Confidence feature selection, fusion, classifier architectures, and training settings are detailed in Supplementary Section S7, with results in Supplementary Table S6.

#### Binding Affinity Prediction

The binding affinity prediction head uses an interface mixture-density network (MDN) [3]. For each inter-entity interface edge (*i, j*) ∈ *ε* _inter_, we form an edge embedding 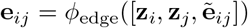, where 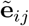 denotes the edge-type embedding. We then use these edge embeddings for the density estimation function. We construct an objective function including three terms: a density loss function [3], a correlation-based function similar to DockQ prediction, and a list-wise ranking loss function [29]. Supplementary Section S3 includes a detailed mathematical description.

#### Residue-level Downstream Evaluation

To assess the quality of OmniScore’s residue-level embeddings, we adopt residue-level prediction tasks of StructTokenBench [17]. In this setting, OmniScore is used as a frozen feature encoder, where StructTokenBench accepts the post-quantization embedding vectors as inputs. For each protein chain, the adapter follows the residue-block convention defined in the Input Preparation section to construct a sequence and a graph. Given a list of quantized embeddings **Z** from a frozen OmniScore, the adapter retrieves an embedding vector 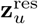 for a residue *u* by performing average pooling on its corresponding atomic embeddings. The resulting matrix 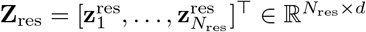 is passed to the internal procedure of StructTokenBench as a continuous input.

### 2.6 Ablation study

We evaluated the contributions of model components using the combinations summarized in Table 3. The Base configurations used a Mamba-based autoencoder with RFSQ, whereas the BaseGeo configurations additionally incorporated EGT before the Mamba encoder. Within these groups, we compared auxiliary property prediction, token and coordinate masking, and residue exposure features and prediction objectives. We also varied model size and quantization settings, including the number of residual quantization layers and the scalar quantization levels. We assessed reconstruction RMSD and downstream StructTokenBench performance using frozen pretrained encoders, and additionally measured code-usage entropy and utilization for the quantization comparisons.

**Table 3:** Model Components and Combinations. Rows present modules in OmniScore, while columns represent different component combinations grouped into two big classes. The Base group contains model variants derived from the standard Mamba-based Autoencoder module, while the BaseGeo group includes ones that extend from the *BaseGeo*, which adds EGT before the Mamba encoder of the autoencoder.

| Component | Base |  |  | BaseGeo |  |  |  |  |  |  |  |  |
| --- | --- | --- | --- | --- | --- | --- | --- | --- | --- | --- | --- | --- |
|  | Base | RTM<br>+ SM | RTM<br>+ SM<br>+ MHP | Base<br>Geo | EP | MHP | RTM | CTM | CTM<br>+ SM | RTM<br>+ SM | REF | REP |
| **Mamba-based Autoencoder** |  |  |  |  |  |  |  |  |  |  |  |  |
| *A foundational model based on Mamba encoder-decoder with RFSQ* | ✓ | ✓ | ✓ | ✓ | ✓ | ✓ | ✓ | ✓ | ✓ | ✓ | ✓ | ✓ |
| **Atom/Block Tokens** |  |  |  |  |  |  |  |  |  |  |  |  |
| *Discrete token embeddings for atoms and residues* | ✓ | ✓ | ✓ | ✓ | ✓ | ✓ | ✓ | ✓ | ✓ | ✓ | ✓ | ✓ |
| **Geometric GNNs (Geo)** |  |  |  |  |  |  |  |  |  |  |  |  |
| *Generalist geometric graph neural networks* |  |  |  | ✓ | ✓ | ✓ | ✓ | ✓ | ✓ | ✓ | ✓ | ✓ |
| **Energy Prediction (EP)** |  |  | ✓ |  | ✓ | ✓ | ✓ | ✓ | ✓ | ✓ | ✓ | ✓ |
| *Rosetta score regression from CLS token* |  |  |  |  |  |  |  |  |  |  |  |  |
| **Multi-Head Prediction (MHP)** |  |  |  |  |  |  |  |  |  |  |  |  |
| *Multiple regression targets (DockQ, K<sub>d</sub>, K<sub>i</sub>, IC<sub>50</sub>)* |  |  | ✓ |  |  | ✓ | ✓ | ✓ | ✓ | ✓ | ✓ | ✓ |
| **Random Token Masking (RTM)** |  | ✓ | ✓ |  |  |  | ✓ |  |  | ✓ | ✓ | ✓ |
| *Independent atom/block token corruption* |  |  |  |  |  |  |  |  |  |  |  |  |
| **Continuous Token Masking (CTM)** |  |  |  |  |  |  |  | ✓ | ✓ |  |  |  |
| *Continuous span token corruption* |  |  |  |  |  |  |  |  |  |  |  |  |
| **Structure Masking (SM)** |  | ✓ | ✓ |  |  |  |  |  | ✓ | ✓ | ✓ | ✓ |
| *3D coordinate corruption during encoding* |  |  |  |  |  |  |  |  |  |  |  |  |
| **Residue Exposure Features (REF)** |  |  |  |  |  |  |  |  |  |  | ✓ | ✓ |
| *SASA/HSE Features: solvent accessibility and burial features* |  |  |  |  |  |  |  |  |  |  |  |  |
| **Residue Exposure Prediction (REP)** |  |  |  |  |  |  |  |  |  |  |  | ✓ |
| *Per-residue surface area prediction* |  |  |  |  |  |  |  |  |  |  |  |  |

## 3 Results

### 3.1 Model Quality and Interaction Assessment

One of the key criteria for a successful 3D biomolecular structure embedding is the ability to distinguish native or native-like structures from non-native ones. Here, we hypothesized that if a given embedding model successfully learned the spatial distribution of individual atoms comprising native biomolecular structures, the embedding vectors of non-native structures would be mapped to relatively sparsely populated regions of the learned native distribution. In addition, since native conformations are closely linked to binding affinities of biomolecular complexes, high-quality structural embeddings may also facilitate accurate affinity predictions, as shown in protein–ligand affinity models [3, 30]. We evaluated OmniScore on two complementary groups of structure-level tasks: model quality assessment of antibody– antigen complexes and RNA structures, and interaction prediction for pMHC and protein–ligand complexes. Together, these tasks assess whether the learned representations capture structural accuracy and binding-related properties across diverse biomolecular systems.

#### Antibody–antigen model quality assessment

On antibody–antigen structure quality assessment [7], OmniScore achieved the highest point estimates on all four comparison metrics among the evaluated baselines (Fig. 2a). We report Pearson and Spearman correlations, the area under the receiver operating characteristic curve (AUROC), and the area under the precision–recall curve (AUPRC). Among OmniScore variants, the one with 111M parameters reached the highest overall scores, with Pearson correlation of 0.909, Spearman correlation of 0.848, AUROC of 0.990, and AUPRC of 0.990, while the remaining variants showed competitive performance across the four metrics (Fig. 2a). Compared to IgPose [7], the strongest baseline evaluated on the dataset, the best OmniScore variant improved Pearson correlation by 0.21, Spearman correlation by 0.14, AUROC by 0.11, and AUPRC by 0.12.

**Figure 2:**
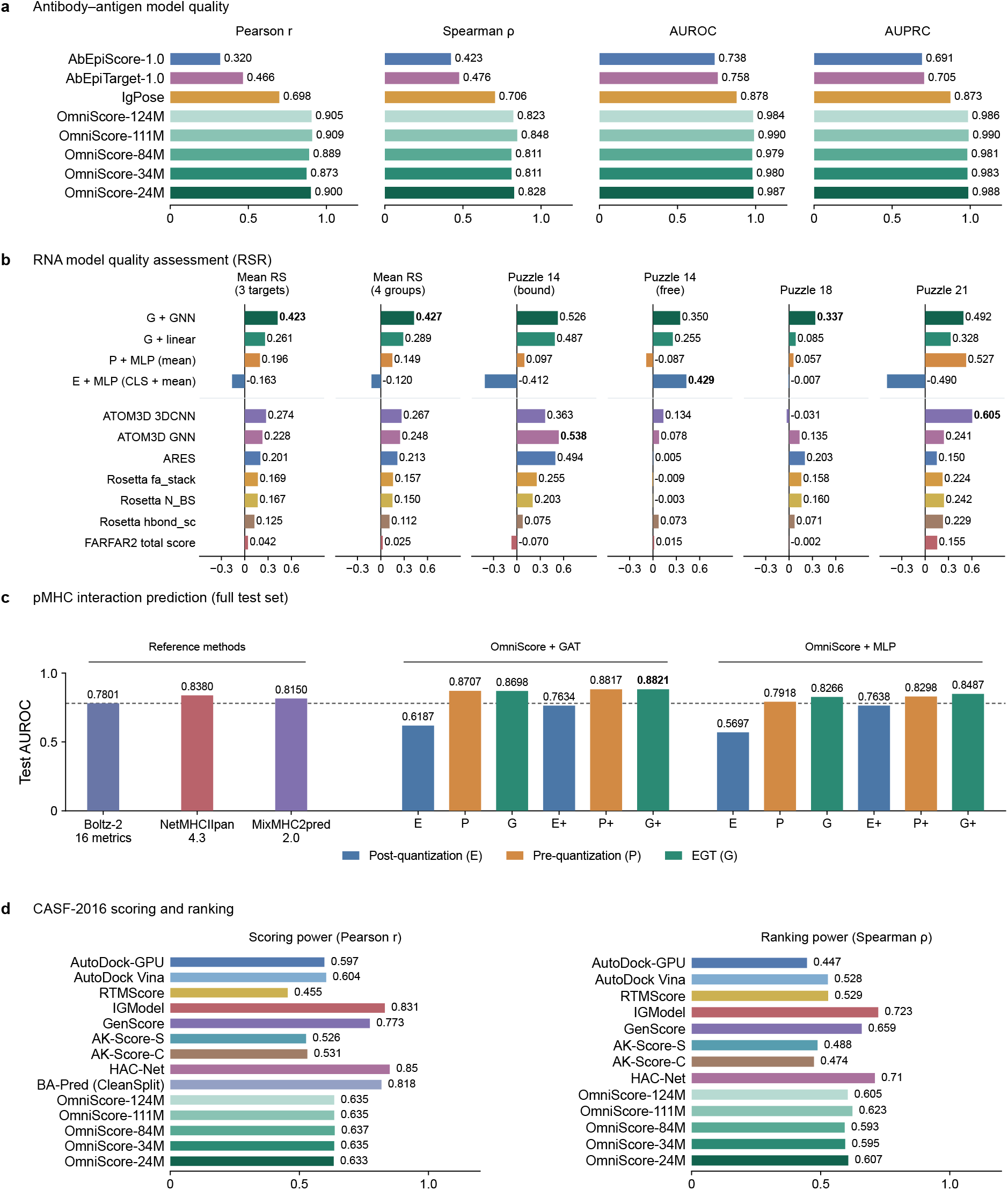
OmniScore performance across four structure-level tasks. **a**, Antibody–antigen model quality on SID-R [7, 10]. **b**, RSR Spearman correlations for RNA Puzzles 14 (bound/free), 18, and 21 after removal of RNA3DB–RNA-Puzzles overlap. The three-target mean first averages the bound/free correlations for Puzzle 14, then averages across the three RNAs; the four-group mean weights all four structure groups equally. Columns share one scale and bold indicates column maxima. Linear denotes the centered EGT probe. ATOM3D baselines were retrained with three-seed ensembles [22]; ARES uses released weights [28]. Full results appear in Supplementary Table S5. **c**, pMHC classification on the full test set of the Qualitative dataset from PREpiBind [25] (Supplementary Table S6). E, P, and G denote post-quantization, pre-quantization, and EGT representations; “+” denotes added Boltz-2 confidence features in panel c. The dashed line marks the confidence-only test AUROC (0.7801). **d**, CASF-2016 scoring and ranking [18]. Literature baselines are from Refs. [31, 32]; BA-Pred (CleanSplit) is shown only for scoring. HAC-Net values use two decimals; others use three. Uncertainty estimates are not shown.

#### RNA structure ranking

We further evaluated OmniScore on the ATOM3D RNA structure ranking task [22] using EGT outputs (G), pre-quantization Mamba outputs (P), and post-quantization embeddings (E). We used mean RS to assess how well predicted RMSD ranks candidates within each RNA target. The evaluation comprised RNA Puzzles 14 (bound/free), 18, and 21, with overlap between RNA3DB and RNA-Puzzles data removed before evaluation. A GNN head using frozen G embeddings achieved a three-target mean RS of 0.423 (four-group mean, 0.427; Fig. 2b; Supplementary Table S5). This exceeded the retrained ATOM3D 3DCNN (0.274) and GNN (0.228) baselines, ARES with released weights (0.201) [28], and the best evaluated individual Rosetta score term, `fa_stack` (0.169). The FARFAR2 Rosetta total score achieved 0.042 [23]. The GNN correlations were positive for all four structure groups (0.526, 0.350, 0.337, and 0.492 for Puzzle 14 bound/free, 18, and 21, respectively).

The centered linear probe on G achieved a mean RS of 0.261, whereas MLP heads using mean-pooled P and CLS plus mean-pooled E embeddings achieved 0.196 and 0.163, respectively. These results support the utility of pretrained graph representations for RNA model assessment, while showing that their benefit depends on the representation and prediction head. The weaker results from later representations under the tested heads do not establish that quantization alone causes a loss of ranking information. Performance varied across the structure groups: for example, the retrained 3DCNN achieved the highest correlation on Puzzle 21 (0.605), despite its lower mean RS. Additional representation and head comparisons are provided in Supplementary Section S6.

#### pMHC interaction prediction

We also evaluated these three representation stages for pMHC interaction classification. OmniScore classifiers were compared with widely used NetMHCIIpan-4.3, MixMHC2pred-2.0, and a baseline prediction model using 16 Boltz-2 confidence metrics (Supplementary Table S6; Fig. 2c).

Combining Boltz-2 confidence metrics with frozen OmniScore embeddings improved test AUROC, supporting the utility of these embeddings for interaction prediction. With GAT classifiers, combining P or G with confidence metrics yielded AUROCs of 0.882 for both P+ and G+, compared with 0.780 for confidence metrics alone. Adding confidence metrics also improved the MLP classifiers, yielding AUROCs of 0.830 and 0.849 for P+ and G+, respectively. On the same full test set, NetMHCIIpan-4.3 and MixMHC2pred-2.0 achieved AUROCs of 0.838 and 0.815, respectively, below the GAT configurations using G alone (0.870) or G+ (0.882). These results suggest that both continuous sequence and graph representations provide information complementary to structure-prediction confidence.

#### Protein-ligand interaction prediction

On CASF-2016 protein–ligand benchmark [18] (Fig. 2d), OmniScore produced similar scoring and ranking results across model sizes, exceeding several evaluated baselines but remaining below the strongest specialized methods. The OmniScore models achieved Pearson’s *r* values of 0.633–0.637 on the scoring power and Spearman’s *ρ* values of 0.593–0.623 on the ranking power benchmarks. Specifically, its variants reached Pearson’s *r* of 0.635 (OmniScore-124M), 0.635 (OmniScore-111M), 0.637 (OmniScore-84M), 0.635 (OmniScore-34M), and 0.633 (OmniScore-24M) for the scoring task. Even though all variants fell below HAC-Net (0.85) [31, 33], IGModel (0.831) [34], BA-Pred (CleanSplit, 0.818) [32], and GenScore (0.773) [3], they exceeded AutoDock Vina (0.604) [35], AutoDock-GPU (0.597) [36], RTMScore (0.455) [37], and both AK-Score-S (0.526) [2] and AK-Score-C (0.531) [2]. For ranking protein–ligand complexes, OmniScore’s variants reached Spearman’s *ρ* of 0.605 (OmniScore-124M), 0.623 (OmniScore-111M), 0.593 (OmniScore-84M), 0.595 (OmniScore-34M), and 0.607 (OmniScore-24M). These results were below IGModel (0.723) and GenScore (0.659), while exceeding AutoDock Vina (0.528), RTMScore (0.529), AutoDock-GPU (0.447), AK-Score-S (0.488), and AK-Score-C (0.474).

### 3.2 Residue-level downstream prediction

On StructTokenBench (Table 4), OmniScore showed competitive overall performance among the evaluated protein-specialized tokenization methods across 15 functional-site classification settings and 6 physicochemical-property regression tasks. OmniScore-84M and OmniScore-34M achieved average AUROC values of 71.76% and 70.90% for functional-site prediction tasks, respectively. They clearly outperformed ESM3, VanillaVQ, ProTokens, and FoldSeek by up to 19.86 percentage points, ranking second and third among the evaluated methods, slightly behind AminoAseed (72.43%). At the task level, both OmniScore variants outperformed AminoAseed on six out of 15 classification tasks. These point estimates indicate competitive functional-site prediction performance; statistical equivalence was not established. They consistently improved performance on both BindBio splits, BindShake, CatBio-Fold, Con-Fold, and Rep-Fold, where the highest gaps were observed in CatBio-Fold (75.14% and 77.24% vs. 65.95%) and BindShake (75.16% and 77.75% vs. 69.61%). Specifically, OmniScore-84M obtained the best results on both BindBio splits, BindShake, and CatBio-Fold (70.18%, 70.60%, 77.75%, and 77.24%), while OmniScore-34M achieved the best result on Con-Fold (61.39%).

**Table 4:**
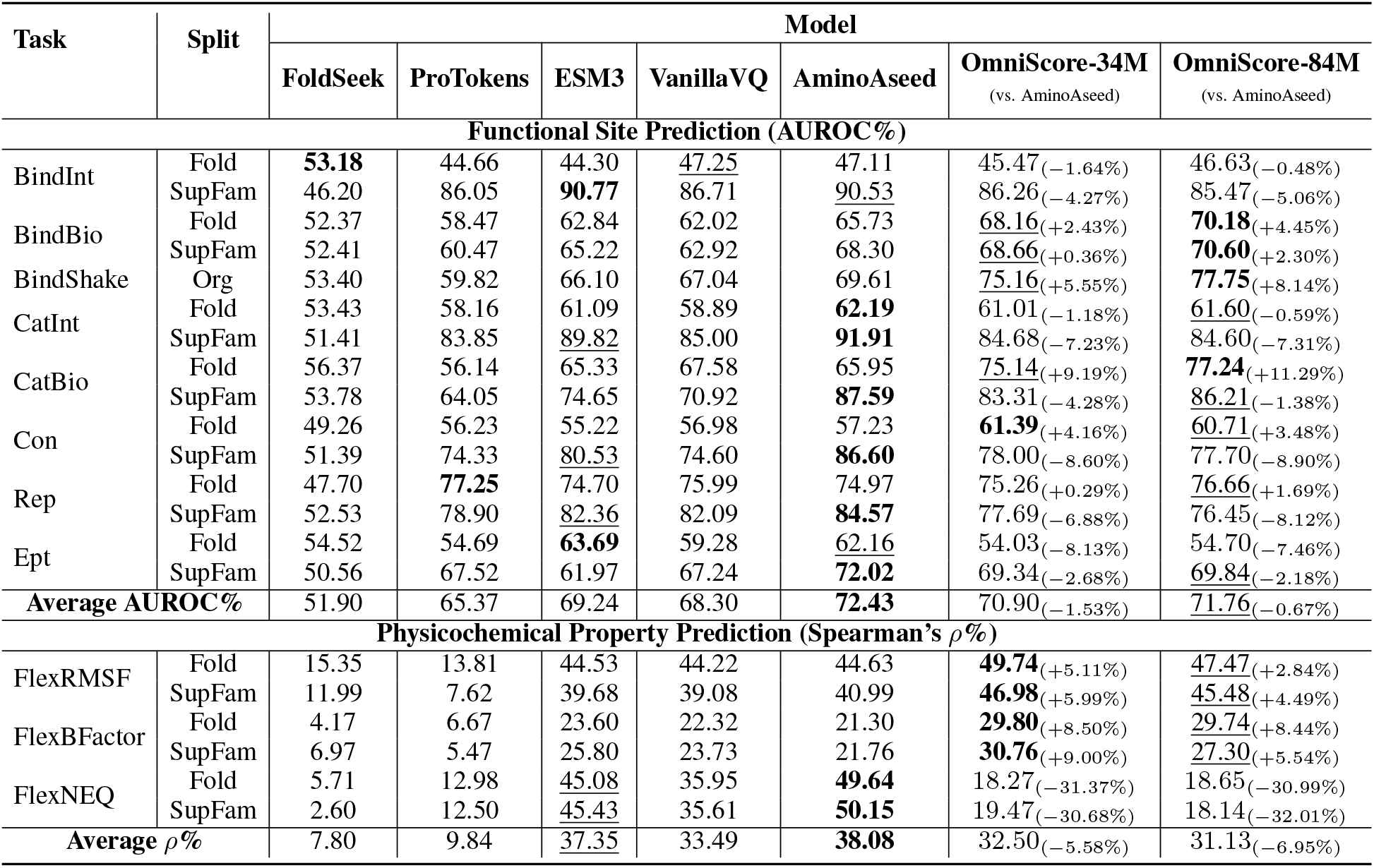
Results on StructTokenBench residue-level benchmark datasets. Bold and underlined text denote the best and second best results in each task. Here, OmniScore variants use a quantization configuration with three layers of RFSQ and levels of [5,5,3,3].

For physicochemical-property regression tasks, the OmniScore-34M and OmniScore-84M models ranked first and second among the tested tokenization methods on the FlexRMSF and FlexBFactor tasks across the Fold and SupFam splits. For the FlexNEQ task, both OmniScore variants showed limited performance, achieving Spearman’s *ρ* values of 18.14–19.47%, which were higher than those of FoldSeek and ProTokens, but considerably lower than those of AminoAseed, ESM3, and VanillaVQ. Overall, these results demonstrate that, despite being trained to cover diverse biological modalities, the OmniScore models achieved performance competitive with that of existing protein-focused models on various protein property prediction downstream tasks.

### 3.3 Reconstruction performance

For biomolecular structure reconstruction, larger models generally achieved better reconstruction performance in the evaluated configurations (Fig. 3a). The best validation reconstruction metrics were obtained with the 1024-dimensional model, OmniScore-111M, achieving an RMSD of 0.689 Å and a template modeling score (TM-score) of 0.998. It is also noticeable that reconstruction performance is not directly linked to downstream task performances. The best downstream classification model in the default-quantizer REP comparison, OmniScore-34M, showed an RMSD value of 2.343 Å and a TM-score value of 0.986 under REP. The best downstream regression model in the same comparison, OmniScore-27M, had an RMSD value of 1.934 Å and a TM-score value of 0.989.

**Figure 3:**
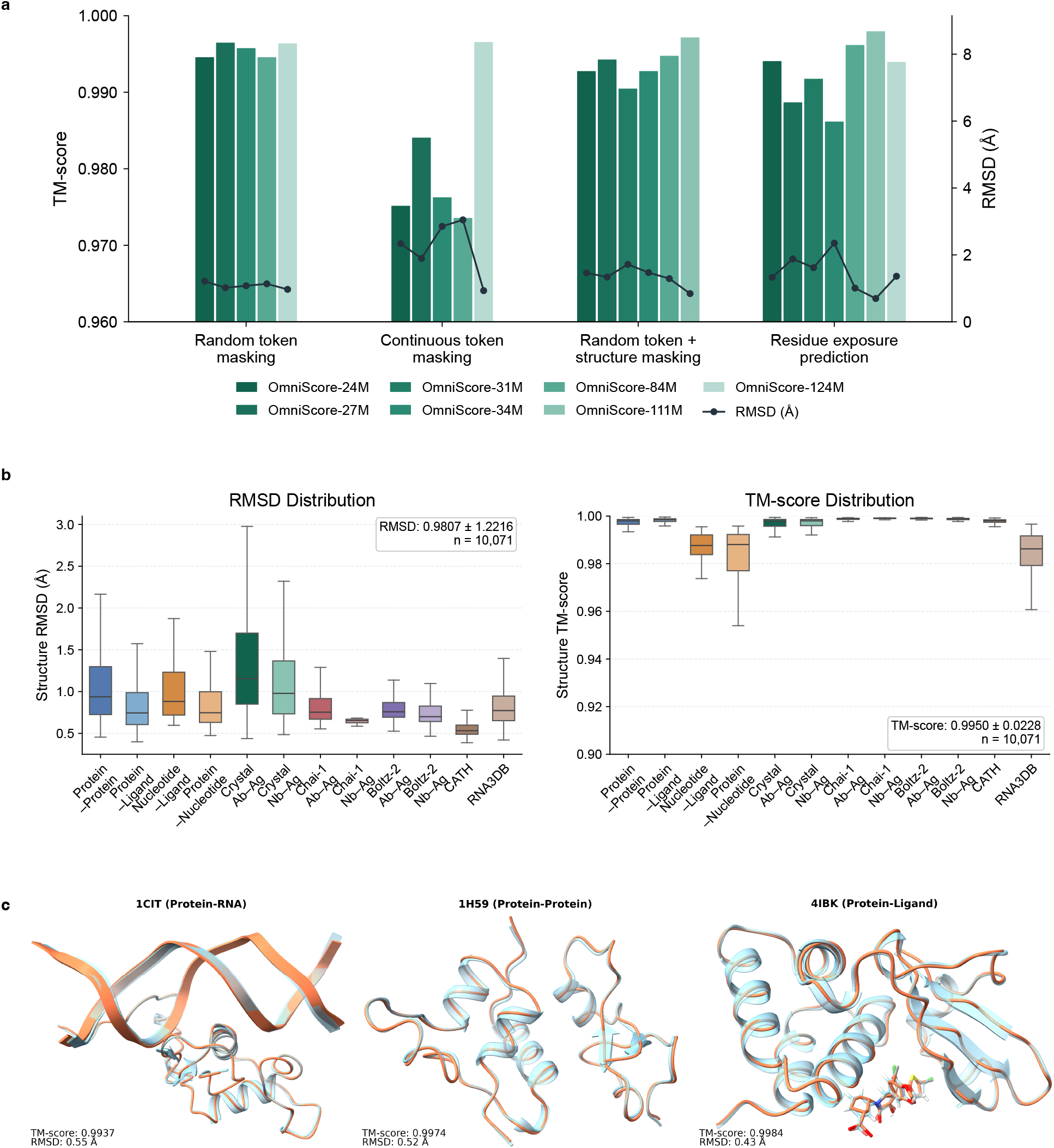
Biomolecular structure reconstruction by OmniScore. **a**, Validation reconstruction performance for seven model sizes (24M–124M parameters) under four pretraining configurations. Bars show TM-score and lines show RMSD. **b**, RMSD and TM-score distributions across validation modalities for OmniScore-84M pretrained with the best setting in Fig. 4. **c**, Representative reconstructions of protein–RNA (1CIT), protein–protein (1H59), and protein–ligand (4IBK) complexes. Reference structures are shown in pale blue and reconstructed structures in orange.

The OmniScore models showed strong overall structure reconstruction accuracy in most tested modalities (Fig. 3b), with an average RMSD value of 0.981 Å and an average TM-score value of 0.995 over 10K structures. In most modalities, TM-score distributions concentrated near 1.0, while the RMSD results varied significantly, with the lowest RMSD for CATH and higher errors for the crystal antibody/nanobody–antigen subsets. The lower TM-score tails were mainly observed for nucleotide–ligand, protein–nucleotide, and RNA3DB subsets, indicating that nucleic-acid-included structures contributed more to the high-error cases than protein-included subsets.

The three qualitative examples (1CIT for protein–RNA, 1H59 for protein–protein, and 4IBK for protein–ligand) further emphasized the aggregate results above (Fig. 3c). All three reconstructed structures approximated the global backbone structures, with TM-score values of 0.994, 0.997, and 0.998 and RMSD values of 0.55 Å, 0.52 Å, and 0.43 Å, respectively. Specifically, the protein regions were well reconstructed in all three examples, with secondary-structure elements and global folds well aligned between prediction and reference. However, even though the protein scaffold looked accurate in the protein–ligand example, the ligand part showed visible local errors, especially around cyclic aromatic groups, suggesting that detailed small-molecule structures remain more difficult to reconstruct than protein backbone geometry.

### 3.4 Ablation Studies

#### Effect of model components

We studied the importance of specific components to both reconstruction performance and downstream tasks of StructTokenBench with three models of different sizes: OmniScore-24M, OmniScore-34M, and OmniScore-84M (Fig. 4, Table 3, Supplementary Table S2 and Supplementary Table S3). The comparison results showed that EGT and multi-head prediction played important roles in structure reconstruction and downstream performance. The Base configuration, a Mamba-based autoencoder only with simple atom/block token embeddings, showed inferior downstream performance across all three model-size groups, with average AUROC values ranging from 64.65% to 65.99%. Only adding masking paradigms (token and structure) and multi-head prediction components to the autoencoder without including EGT did not boost the overall performance, as the strongest models only reached average AUROCs of 64.96–65.99% across the three model-size groups. The reconstruction results showed the same limitation of the Base configurations, where the mean RMSD values remained high, ranging from 3.98 to 21.92 Å.

**Figure 4:**
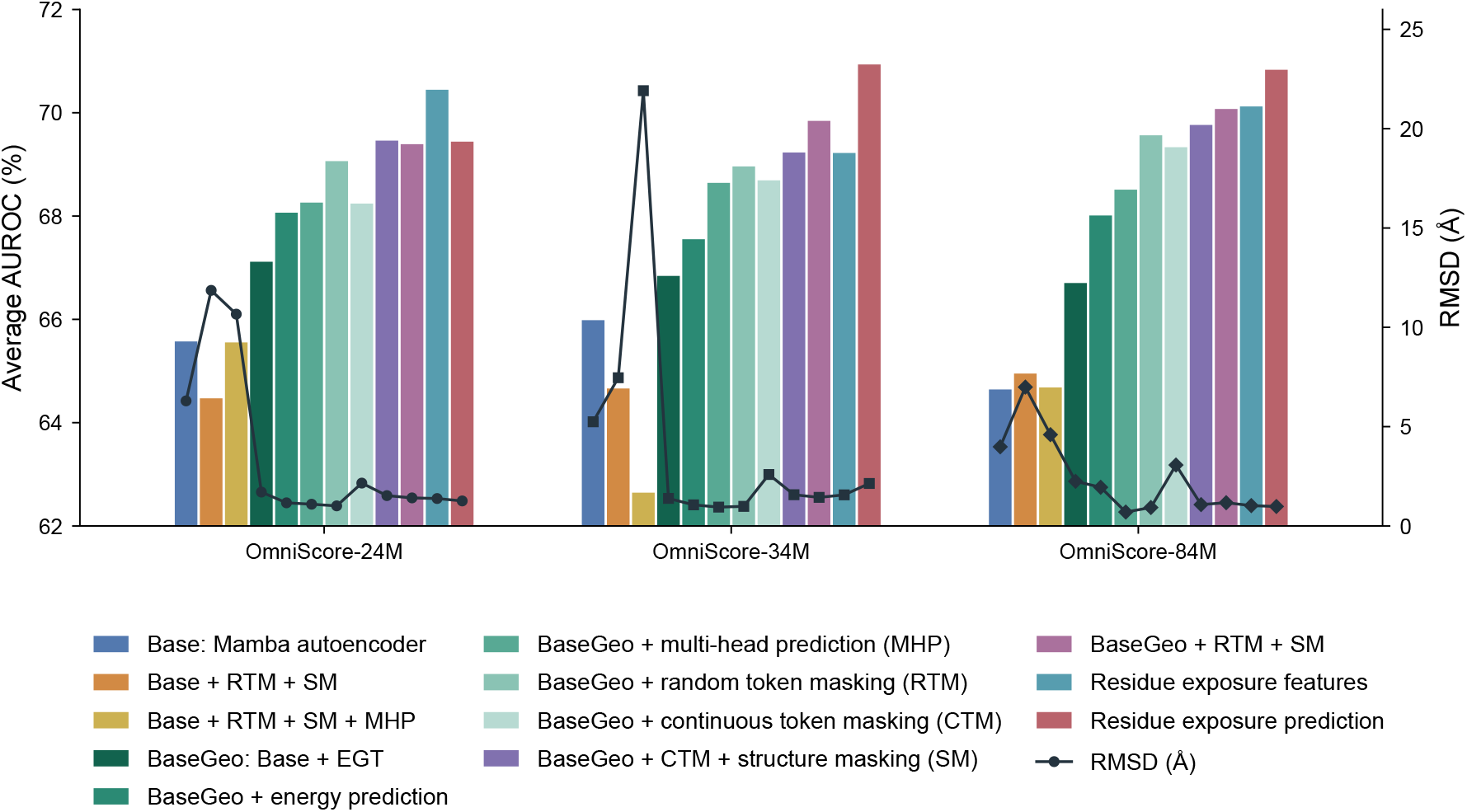
Performance of different training paradigms on the functional-site prediction tasks of StructTokenBench. Each configuration enables different modules and training objectives. In summary, combining EGT with Mamba in the encoder improves average AUROC and reduces RMSD of reconstructed structures. Next, using structure-level information like energy, binding affinity, DockQ and appropriate masking paradigms (coordinates and tokens) can improve the performance further. Finally, residue-level ground-truth is also beneficial to the pretrained models.

In contrast, introducing EGT features into the Mamba autoencoder backbone (BaseGeo) increased the average AUROC to 66.71–67.12%, corresponding to gains of 0.86–2.06 percentage points over the corresponding Base models, and also improved reconstruction quality, with mean RMSDs of approximately 1.39–2.25 Å across the three BaseGeo models. Adding structure-level prediction objectives further improved the BaseGeo backbone. Compared to BaseGeo, the addition of Rosetta energy prediction (EP) (BaseGeo + EP) increased the average AUROC from 66.71% to 68.02% for OmniScore-84M and from 67.12% to 68.07% for OmniScore-24M. Furthermore, adding multi-head prediction for multiple regression targets (BaseGeo + Multi-Head Prediction) produced additional gains; all models exceeded AUROC values of 68%. Reconstruction quality also improved across all three model sizes when adding an energy prediction head and improved further with multiple prediction heads.

Random token masking consistently improved residue-level downstream performance over prediction-head-only models. Specifically, applying random masking to atom tokens after EGT and before the Mamba encoder (BaseGeo + RTM + MHP) increased the average AUROC to above 69% for OmniScore-84M and OmniScore-24M. The 3D coordinate corruption method further improved the masked models, as combining random token masking with structure masking (BaseGeo + RTM + Structure Mask) reached AUROC values of 69.4–70.08%.

Finally, the variants (REF and REP) that included residue exposure information (SASA/HSE) as features and losses led to the highest downstream performance, although the effect slightly varied across model sizes. Particularly, the REF configuration, concatenating these features with EGT features and token embeddings, improved the average AUROC to 70.13% for OmniScore-84M and 70.45% for OmniScore-24M. Adding the auxiliary residue exposure prediction head (REP setting) further increased the AUROC to 70.84% for OmniScore-84M and 70.94% for OmniScore-34M.

#### Model size analysis

Based on the analysis of model components, we analyzed the variation in performance of different model sizes in residue-level downstream tasks by changing the hidden dimension and the number of Mamba encoder and decoder layers under four BaseGeo training configurations (Supplementary Table S2, Supplementary Table S3, Fig. 5). On StructTokenBench, downstream performance did not consistently improve with model size, unlike the general trend in reconstruction performance. With the random token masking setup, OmniScore-84M achieved the best classification result, reaching 69.57% average AUROC, while the best 512-dimensional model, OmniScore-24M, reached 69.07%. In contrast, the 512-dimensional variant, OmniScore-34M, achieved 28.10% average Spearman’s *ρ* in regression tasks, compared with 27.21% for OmniScore-84M.

**Figure 5:**
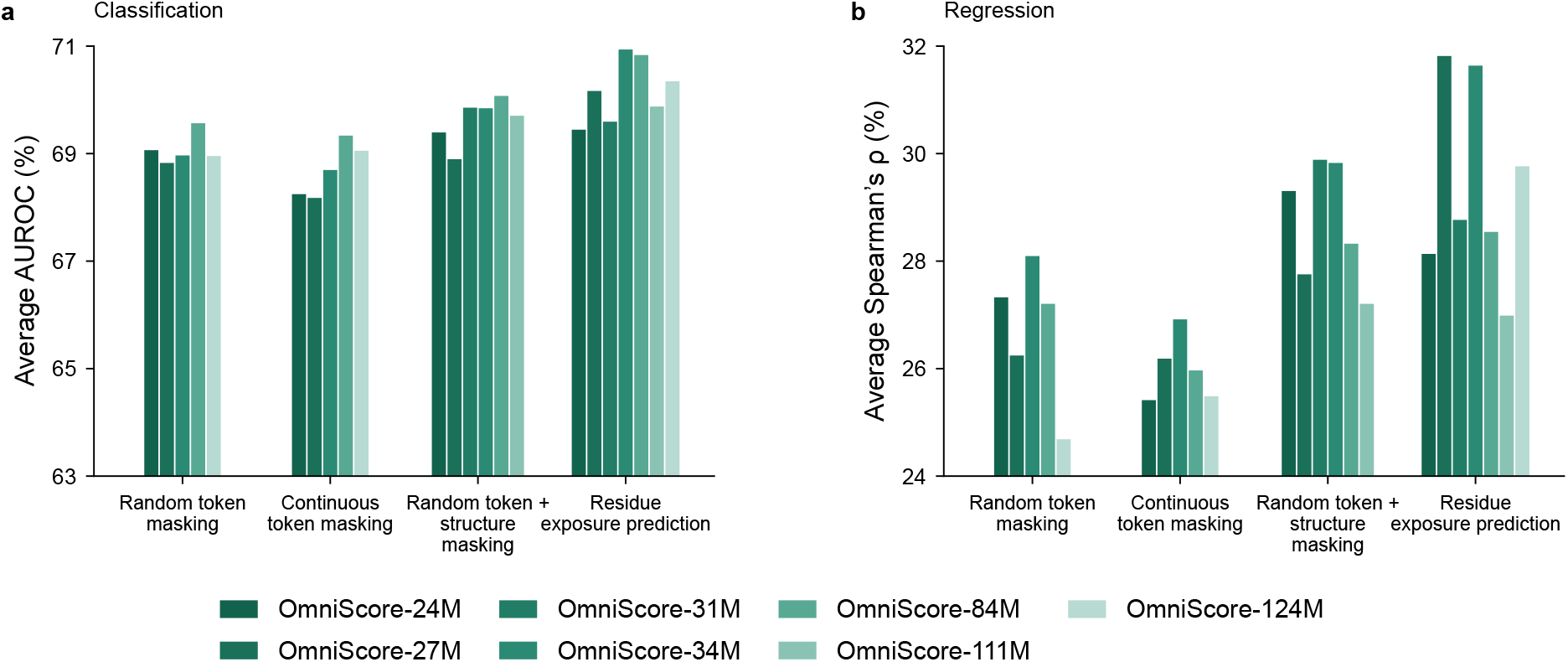
Performance on StructTokenBench of different OmniScore model sizes. We varied the hidden dimensions and the number of Mamba encoder/decoder layers, resulting in model variants 24M, 27M, 31M, 34M, 84M, 111M, and 124M.

After adding 3D coordinate corruption (the last two groups in Fig. 5), the performance difference across models became smaller for classification tasks but remained more visible for regression ones. OmniScore-84M achieved an average AUROC of 70.08%, only 0.22 percentage points above the best 512-dimensional model, OmniScore-31M. In regression, OmniScore-31M reached 29.89% average Spearman’s *ρ*, exceeding OmniScore-84M by 1.56 percentage points. When adding residue exposure features and loss, 512-dimensional models yielded the highest averaged results for both task groups. OmniScore-34M achieved the best classification average at 70.94%, slightly above OmniScore-84M at 70.84%. For regression, OmniScore-27M achieved 31.82%, followed by OmniScore-34M at 31.64%, while the best 1024-dimensional model, OmniScore-124M, reached 29.77%.

#### Quantization configuration

To assess the tokenizer as a component of encoder pretraining, we compared six quantization settings across four OmniScore sizes using the residue exposure prediction (REP) configuration in Fig. 4: single-layer FSQ with levels 4^6^ or 5^8^, two-layer RFSQ with levels [7, 5, 5, 5] or [8, 5, 5, 3], three-layer RFSQ with levels [5, 5, 3, 3], and four-layer RFSQ with levels [4, 3, 2, 2] (Fig. 6).

**Figure 6:**
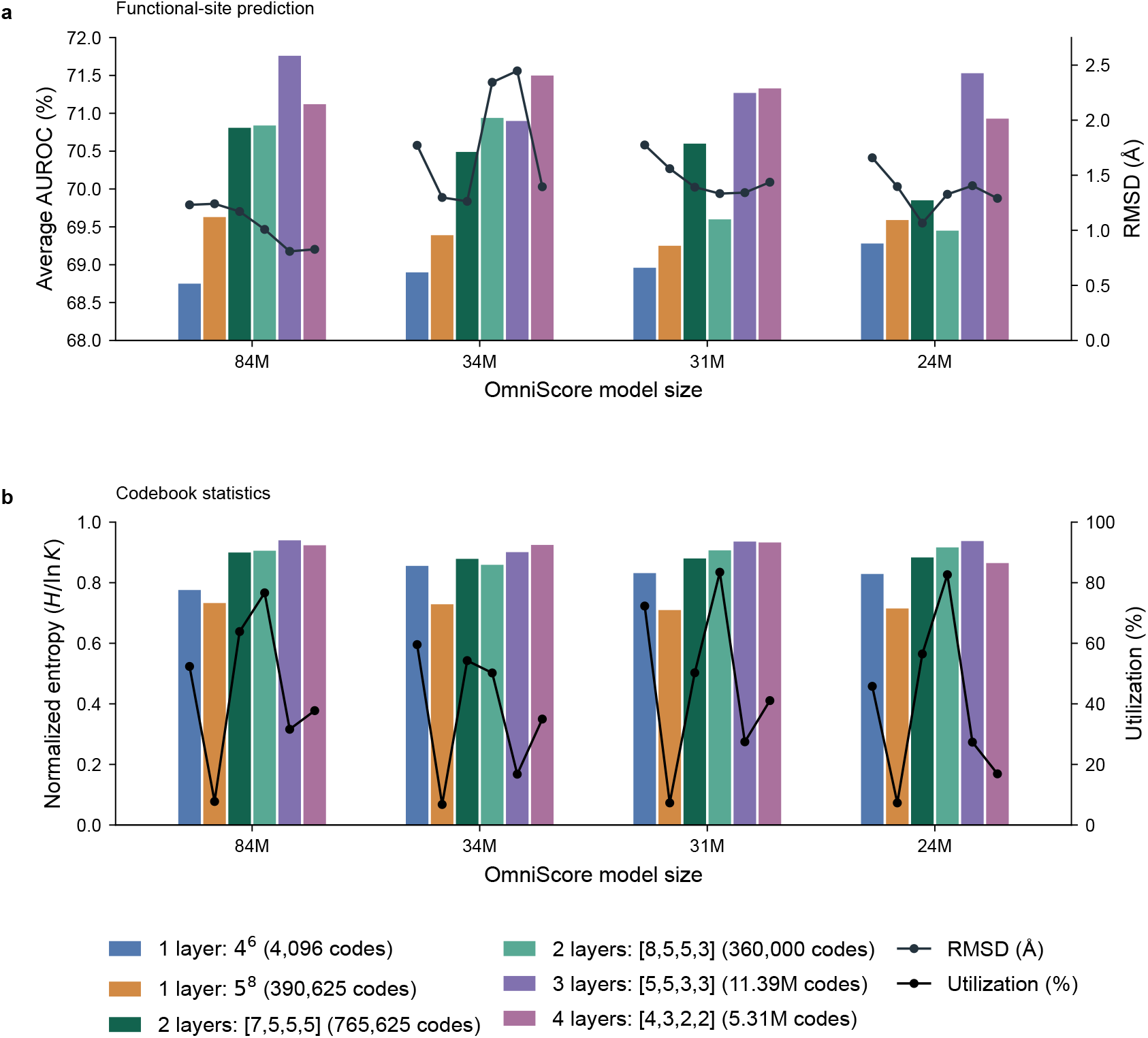
Quantization configurations across the four model groups. Top: StructTokenBench performance, where bars show the average AUROC (left axis) and the line shows RMSD (right axis). Bottom: codebook usage, where bars show the normalized entropy *H/* ln *K* (left axis) and the line shows utilization (right axis). Both panels share the six-color quantizer legend, and the two lines use separate colors.

Residual quantization improved functional-site prediction over single-layer FSQ, which achieved average AUROCs of 68.75–69.63%. Three-layer RFSQ performed best for OmniScore-84M (71.76%) and OmniScore-24M (71.53%), whereas four-layer RFSQ performed best for OmniScore-34M (71.50%) and OmniScore-31M (71.33%). Reconstruction gains depended on model size: for OmniScore-84M, three-layer RFSQ reduced RMSD from 1.23–1.24 Å to 0.80 Å, but smaller models showed no consistent improvement with residual depth. We used three-layer RFSQ with levels [5, 5, 3, 3] for the StructTokenBench comparisons in Table 4.

Code-usage statistics demonstrate that higher utilization did not necessarily correspond to better downstream performance, consistent with the broader distinction between reconstruction quality and task-specific representation utility. Normalized entropy (*H/* ln *K*) measures the uniformity of code assignments, where *H* is usage entropy and *K* is codebook size; utilization measures the fraction of codes observed on the validation set. Single-layer FSQ with levels 5^8^ had the lowest entropy (0.710–0.733) and utilization (6.8–7.8%). Three-layer RFSQ achieved higher entropy (0.901–0.940), although only 16.8–31.6% of its approximately 11.39 million codes were used. Two-layer RFSQ with levels [8, 5, 5, 3] yielded utilization of 50.2–83.5%, reaching the highest observed value across all configurations. Because residual depth and code-space capacity varied together, these comparisons did not isolate the effect of residual depth alone.

## 4 Discussion

Traditional model quality and affinity scoring approaches remain largely fragmented, relying on specialized encoders designed for specific modalities. This necessity for distinct architectures across different tasks has hindered scalable modeling of complex, multi-component systems. In contrast, our evaluations support OmniScore as a shared pretrained geometric encoder for biomolecular representation learning, with tokenization serving as one component of the pretraining framework. Specifically, OmniScore outperformed the evaluated baselines [7, 10] on the antibody–antigen model quality assessment task [7]. The RNA results similarly showed the utility of graph representations for candidate ranking. The pMHC results further indicate that the utility of OmniScore embeddings depends on the representation stage and adaptation strategy. G+ achieved the highest observed AUROC among the frozen-backbone configurations, closely followed by P+; GAT classifiers generally outperformed MLP classifiers. On StructTokenBench [17], OmniScore approached the strongest evaluated tokenization baseline in average functional-site AUROC and led four flexibility-related regression settings, while retaining substantial limitations on FlexNEQ. On the CASF-2016 protein–ligand benchmark [18], OmniScore exceeded several evaluated scoring functions but remained below the strongest specialized methods on both scoring and ranking power.

OmniScore connects specialized biomolecular scoring [1, 2, 3, 5, 7, 10, 34], geometric representation learning [11, 12, 38], and structural tokenization [17, 21] through a shared pretrained encoder. Within this broader landscape, OmniScore’s contribution centers on the reuse of frozen geometric representations across molecular classes and prediction scales, from complex-level model quality assessment and interaction prediction to residue-level functional and flexibility prediction. Within this framework, Bio2Token [21] provides a closely related Mamba-based autoencoder architecture, while OmniScore incorporates geometric graph encoding and structural and hysicochemical supervision to support downstream representation learning. BioScore [12] shares the goal of assessing diverse biomolecular complexes through an interface-based scoring potential; OmniScore provides representations that can be adapted through task-specific prediction heads. The evaluations further distinguish the utility of graph, continuous sequence, and quantized representations, showing that the preferred representation depends on the task and that reconstruction quality alone does not reliably indicate downstream performance.

Ablation studies show that our design choices, using EGT, auxiliary losses, and residue exposure information, helped improve representation learning. The weak performance of the *Base* model, which uses Mamba for both the encoder and decoder as in [21], suggests that sequence modeling alone was insufficient to capture the geometric information needed for the evaluated tasks under this pretraining setup. Compared to the *Base* model, the *BaseGeo* model with EGT before Mamba led to higher accuracy in both reconstruction and downstream performances. Furthermore, introducing additional auxiliary losses, such as physics-based energy, DockQ, or binding affinity, and per-residue exposure features improved downstream performance in several configurations. These results suggest that structural and physicochemical supervision is a useful strategy for learning transferable biomolecular representations.

During pretraining, reconstruction quality provided a useful training signal but was not a reliable indicator of downstream performance. Specifically, within the default-quantizer REP comparison, our larger 1024-dimensional models produced the best validation reconstruction, yet the smaller 512-dimensional models gave the strongest StructTokenBench classification and regression results. This mismatch indicates that selecting models solely by RMSD or TM-score may favor representations that are less effective for downstream prediction, because a latent space that reproduces coordinates faithfully may not preserve the information most relevant to downstream prediction. A related limitation emerged in the subset and case-study examples: global folds were recovered well, while ligand ring geometry and nucleic-acid-containing systems remained difficult. These observations motivate combining reconstruction metrics with lightweight downstream probes for model and checkpoint selection, while evaluating reconstruction through both global fold recovery and local chemical fidelity.

Based on these results and analyses, we can expand and improve OmniScore further in several directions. In the current version, we mainly focused on stabilizing the pretraining process and benchmarking residue-level embeddings, while implementation of the prediction heads is not extensively optimized. Thus, further improvement of the prediction heads for specific downstream tasks will make them more useful and accurate for practical usage. The current pretrained models also suffered from high reconstruction losses and low visualized reconstruction quality with nucleic-acid-containing complexes and small molecules, potentially reflecting limited data and incomplete geometric supervision. Although relative scores can help to rank and sort protein–ligand complexes, a curated absolute scoring function will make the output scores more meaningful and intuitive in practical scenarios. Broader external evaluations with explicit overlap checks between pretraining and test data, repeated runs, and target-level uncertainty estimates are needed to establish robustness, particularly for RNA ranking with only three independent test targets represented by four structure groups. Matched pretraining controls using graph-only encoders or continuous latents without quantization would help isolate the contributions of sequence modeling and tokenization. More extensive optimization of the combinations of model components can be performed for further improvement.

Recently, new drug modalities are actively developed by conjugating multiple modalities such as antibody–drug, antibody–oligonucleotide, and peptide-drug conjugates to treat previously untreatable diseases [39]. OmniScore’s shared geometric representation framework provides a foundation for extending structure-based modeling and prediction to such multi-component systems and may facilitate drug discovery and design of new modalities.

## 5 Conclusion

Assessing the quality of biomolecular complexes remains difficult because proteins, nucleic acids, small molecules, and their interfaces differ in size, chemistry, and available supervision, and most existing methods address only one molecular class at a time. This fragmentation limits how well a single scoring function can transfer across interaction types as structure prediction models produce increasingly diverse complexes. We introduced OmniScore to address this problem through a shared pretrained geometric encoder whose frozen representations are reused through task-specific prediction heads.

OmniScore learns a shared geometry-aware latent space of diverse types of biomolecules: proteins, nucleic acids, and small molecules. It converts each structure into two coupled views, a block-level heterogeneous graph and an aligned atom-wise sequence, and encodes them with an equivariant graph transformer followed by a bidirectional Mamba encoder. An RFSQ tokenizer maps the continuous embeddings into compact latent representations as one component of the pretraining framework. The model is pretrained on both complexes and monomers through coordinate reconstruction, corruption recovery, atom and block type classification, and structure-level regression. The pretrained encoder provides graph, continuous sequence, and quantized representations for task-specific heads during downstream adaptation.

OmniScore outperformed the evaluated baselines on antibody–antigen quality assessment. Graph representations also supported RNA structure ranking, where a GNN head exceeded the retrained ATOM3D baselines after removal of RNA3DB–RNA-Puzzles overlap. For pMHC interaction classification, combining OmniScore embedding with Boltz-2 confidence features yielded high prediction accuracy supporting the utility of the learned geometric representations. On CASF-2016, OmniScore exceeded several baselines but remained below the strongest specialized methods. For downstream tasks, OmniScore residue embeddings approached the strongest evaluated baseline in average functional-site AUROC and led four flexibility-related regression settings. The ablation results indicate that combining geometric graph features with sequential modeling improved both reconstruction performance and downstream transfer. OmniScore provides a common basis for developing structure-based prediction models across molecular classes, with the tokenizer supporting the broader goal of transferable geometric representation learning. The shared representation of OmniScore will serve as a foundation for developing future computational methods for emerging multi-component therapeutics.

## 6 Acknowledgements

This work was supported by grants from the Korea Evaluation Institute of Industrial Technology (KEIT) funded by the Ministry of Trade, Industry and Resources (MOTIR), Republic of Korea (Project Nos. RS-2025-25460612 and RS-2026-25617196), and by the Bio & Medical Technology Development Program of the National Research Foundation of Korea (NRF) funded by the Korean government (MSIT) (Project No. RS-2025-25396539). This work was also supported by the Korea Bio Data Station (K-BDS), which provided computing resources and technical support. This research was supported by the “Advanced GPU Utilization Support Program” funded by the Government of the Republic of Korea (Ministry of Science and ICT) (No. 04-26-03-0014).

## Notes

### Competing Interest Statement

The authors have declared no competing interest.

### Summary of Updates

Additional experimental results have been updated.

