## Supplementary Material v2 for "OmniScore: A geometric foundation encoder for universal biomolecular representation and scoring"

### S1 Architecture notation and model configurations

#### S1.1 Architecture notation

Table S1: Essential notations used to describe OmniScore architecture.

| Symbol | Meaning |
|---|---|
| $S$ | Ordered, flattened atom-wise sequence |
| $L$ | Number of atom rows in $S$ |
| $\mathcal{B}$ | Block-type set $\{\text{residue}, \text{nucleotide}, \text{atom}\}$ |
| $a_i$ | Atom token for row $i$, such as carbon, nitrogen, oxygen, or a special token |
| $b_i$ | Block token for row $i$, such as residue, nucleotide, or small molecular atom identity |
| $t_i$ | Block type for row $i$ |
| $\mathbf{x}_i$ | Coordinate of atom $i$ |
| $\mathbf{u}_i$ | Input vector for atom row $i$ to the Mamba encoder |
| $d$ | Hidden dimension of the continuous and quantized Mamba embeddings |
| $d_a, d_b, d_G$ | Dimensions of the atom-token, block-token, and EGT embeddings, respectively |
| $c$ | Number of predefined atom channels per block, varying by block type |
| $\delta(\mathbf{x}_i, \mathbf{x}_j)$ | The distance between atom coordinates $\mathbf{x}_i$ and $\mathbf{x}_j$ in 3D space |
| $\mathcal{G} = (\mathcal{V}, \mathcal{E}, T, R, \mathbf{X})$ | Heterogeneous geometric graph |
| $\mathcal{V}$ | Graph nodes, where each node is a block |
| $\mathcal{E}$ | Typed block-to-block graph edges |
| $T$ | Atom and block token attributes |
| $R$ | Block-type and edge-type attributes |
| $\mathbf{X}$ | Target coordinate matrix |
| $\hat{\mathbf{X}}$ | Decoded/reconstructed coordinate matrix |
| $N$ | Number of blocks in $\mathcal{G}$ |
| $\mathbf{H}^G$ | Equivariant graph transformer's output after graph encoding |
| $\mathbf{H}$ | Continuous sequence embedding before quantization |
| $\mathbf{Z}$ | Quantized decoder latent sequence |
| $\mathbf{z}_{\text{CLS}}$ | Global classification (CLS) representation used for sequence-level heads |
| $\Omega_{\text{atom}}$ | Set of atoms comprising the input molecular structure, indexed by their positions in the flattened sequence; excludes special tokens and padding |
| $\mathcal{M}$ | Set of selected atom-row positions for token masking |

#### S1.2 Model configurations

Table S2: OmniScore model variants with different hidden dimensions and Mamba encoder/decoder depths. Parameter counts by component are reported in Supplementary Table S3.

| Model Name | Hidden Dimension | Mamba Encoder/Decoder Depth |
|---|---|---|
| OmniScore-124M | 1024 | 12 / 6 |
| OmniScore-111M | 1024 | 12 / 4 |
| OmniScore-84M | 1024 | 8 / 4 |
| OmniScore-34M | 512 | 12 / 6 |
| OmniScore-31M | 512 | 12 / 4 |
| OmniScore-27M | 512 | 8 / 6 |
| OmniScore-24M | 512 | 8 / 4 |

#### S2 Graph Construction

As the number of atoms in residues and nucleotides varies, we normalize the block design so that each block stores a fixed number of atom channels for its block type. This design choice aligns tensors within each block type, accelerates batching, and makes the implementation easy to update when new block types or atom layouts are added. For each block $u \in \mathcal{V}$, the number of atom channels $c$ is set to 4 for residues, 9 for nucleotides, and 1 for individual atoms.

Residue blocks represent residues with N, CA, C, and the last side-chain carbon in the atom-row sequence. Glycine has no side-chain carbon, so CA is reused as its fourth residue channel. If the last side-chain carbon is missing for any other residue, CA is also used for that channel. This choice keeps backbone geometry and one side-chain direction while controlling the sequence length. Nucleotide blocks use P, O5', C5', C4', C3', O3', C1', C4, and C6. Atomic blocks support heavy atoms and use one atom per block.

For brevity, graph construction uses undirected edges, where each edge is an unordered pair $\{u, v\}$ associated with an edge type. Edges are built from Euclidean distances $\delta(u, v)$ between block centroids and optional sequence adjacency. Specifically, for each intra- or inter-entity relation $r$, we first retain node pairs whose distance does not exceed a relation-specific cutoff $\tau_r$. We then apply symmetric $k$-nearest-neighbor (KNN) filtering, retaining a pair whenever either node belongs to the other node's $k$ nearest candidates ($k = 10$ in practice). This procedure yields the intra-entity and inter-entity edge sets $\mathcal{E}_{\text{intra}}$ and $\mathcal{E}_{\text{inter}}$, respectively. In practice, we use $\tau_r = 5.0$ Å for intra-entity protein and nucleotide blocks, $\tau_r = 2.0$ Å for covalent neighborhoods between atomic blocks, and $\tau_r = 10.0$ Å for inter-entity neighborhoods. Chain-like protein and nucleotide structures additionally include sequential edges $\mathcal{E}_{\text{seq}}$ connecting adjacent blocks according to their order in the PDB file.

#### S3 Interface MDN Scoring Function

Following the distance-density formulation of GenScore [3], the binding-affinity head assigns a Gaussian mixture to each inter-entity interface edge $(i, j)$. Let $\mathbf{z}_i$ and $\mathbf{z}_j$ be the frozen OmniScore embeddings of its endpoints (e.g. residues), $\tilde{\mathbf{e}}_{ij}$ the frozen edge-type embedding, $\mathbf{e}_{ij} = \phi_{\text{edge}}([\mathbf{z}_i, \mathbf{z}_j, \tilde{\mathbf{e}}_{ij}])$ the edge embedding, and $d_{ij} = \|\mathbf{x}_i - \mathbf{x}_j\|_2$ the observed endpoint distance. The MDN prediction head $g_{\text{MDN}}$, an MLP network with two layers, predicts $K = 10$ mixture weights, means, and standard deviations from $\mathbf{e}_{ij}$ as follows:

$$(\pi_{ij}^{(k)}, \mu_{ij}^{(k)}, \sigma_{ij}^{(k)})_{k=1}^{K} = g_{\text{MDN}}(\mathbf{e}_{ij}), \qquad \sum_{k=1}^{K} \pi_{ij}^{(k)} = 1, \quad \pi_{ij}^{(k)} \geq 0, \quad \sigma_{ij}^{(k)} > 0, \tag{1}$$

The conditional distance density for that edge, denoted by $p_{ij} = p(d_{ij} \mid \mathbf{e}_{ij})$, is

$$p(d_{ij} \mid \mathbf{e}_{ij}) = \sum_{k=1}^{K} \pi_{ij}^{(k)} \frac{1}{\sqrt{2\pi}\, \sigma_{ij}^{(k)}} \exp\left[-\frac{(d_{ij} - \mu_{ij}^{(k)})^2}{2(\sigma_{ij}^{(k)})^2}\right]. \tag{2}$$

OmniScore supports probability-sum and negative log-likelihood (NLL) scoring functions based on the estimated density distributions as follows:

$$\begin{aligned} s_{\text{MDN}}^{\text{prob}} &= \sum_{(i,j) \in \mathcal{E}_{\text{inter}}} p_{ij}, \\ s_{\text{MDN}}^{\text{NLL}} &= -\sum_{(i,j) \in \mathcal{E}_{\text{inter}}} \log p_{ij}. \end{aligned} \tag{3}$$

Based on these functions, we combine a density term with a correlation term and a listwise ranking loss [29] to define the finetuning objective function for the binding-affinity prediction head as follows:

$$\mathcal{L}_{\text{BindingAffinity}} = -\frac{1}{|\mathcal{E}_{\text{inter}}|} \sum_{(i,j) \in \mathcal{E}_{\text{inter}}} \log p_{ij} + 0.7 \times \big(1 - r(s_{\text{MDN}}, y_{\text{pK}})\big) - 0.3 \times \tilde{y}_{\text{pK}} \log \tilde{s}_{\text{MDN}} \tag{4}$$

where $r$ denotes Pearson correlation, $s_{\text{MDN}}$ is the complex-level MDN score and $y_{\text{pK}}$ is the negative-log normalized value of a binding-affinity target, while $\tilde{s}_{\text{MDN}}$ and $\tilde{y}_{\text{pK}}$ are their corresponding batch-wise softmax-based normalized versions. By default, we use $s_{\text{MDN}}^{\text{prob}}$ as the scoring function.

#### S4 Implementation

```
def pretrain(sample):
    G, S, X = construct_graph_and_sequence(sample)

    H_G = f_EGT(G)
    U = concatenate_coordinates_tokens_and_graph_features(
        S, H_G
    )
    U_tilde = optional_dynamic_masking(U)
    H = f_MambaEnc(add_left_right_cls(U_tilde))
    Z_all = q_RFSQ(H)

    z_CLS = Z_all[left_cls] + Z_all[right_cls]
    Z = remove_cls(Z_all)
    y_hat = auxiliary_heads(z_CLS)
    mask_atom_logits, mask_block_logits = mask_token_heads(
        remove_cls(Z_all)
    )
    EGT_atom_logits, EGT_block_logits = EGT_token_heads(H_G)
    X_hat = f_decode(Z)

    return calculate_losses_and_update(
        X_hat, X, y_hat,
        mask_atom_logits, mask_block_logits,
        EGT_atom_logits, EGT_block_logits,
    )
```

Table S3: Model size variants. We use the model size to represent the OmniScore model family, where M means millions of parameters. The character # is the abbreviation symbol of "the number of."

| Model Size | Hidden Dim | # Mamba Encoders | # Mamba Decoders | Total Params | Encoder Mamba | Decoder Mamba | Graph Encoder | Regression Heads |
|---|---|---|---|---|---|---|---|---|
| OmniScore-124M | 1024 | 12 | 6 | 124.8M | 80.4M | 40.0M | 3.0M | 1.3M |
| OmniScore-111M | 1024 | 12 | 4 | 111.5M | 80.4M | 26.7M | 3.0M | 1.3M |
| OmniScore-84M | 1024 | 8 | 4 | 84.8M | 53.7M | 26.7M | 3.0M | 1.3M |
| OmniScore-34M | 512 | 12 | 6 | 34.4M | 20.5M | 10.2M | 3.0M | 657.9K |
| OmniScore-31M | 512 | 12 | 4 | 31.0M | 20.5M | 6.8M | 3.0M | 657.9K |
| OmniScore-27M | 512 | 8 | 6 | 27.6M | 13.8M | 10.2M | 3.0M | 657.9K |
| OmniScore-24M | 512 | 8 | 4 | 24.2M | 13.8M | 6.8M | 3.0M | 657.9K |

##### S4.1 Software Stack

We implemented OmniScore in Python 3.12, PyTorch 2.7, and PyTorch Geometric 2.7. The bidirectional Mamba encoder and decoder used `mamba-ssm` 2.2.5 state-space kernels. We adopted the implementation of the RFSQ algorithm from [40]. We used Biopython v1.87 for structure parsing and feature preparation, RDKit for per-atom chemistry features, and `freesasa` for solvent-accessible-surface-area features. Training and distributed data parallelism were handled by PyTorch Lightning v2.6.

##### S4.2 Default Experimental Configuration

Unless stated otherwise, every experiment shared the following configuration. We used the Adam optimizer with a learning rate of $5 \times 10^{-5}$ without weight decay, under a polynomial schedule of power $3$ in $500{,}000$ iterations, with gradients clipped to a global norm of $1.0$. The quantizer module was fixed to RFSQ with two FSQ layers, each having scalar levels $[8, 5, 5, 3]$. Graph construction used intra-entity cutoffs of $5.0$ Å for proteins and nucleotides and $2.0$ Å for covalent atom–atom edges, an inter-entity cutoff of $10.0$ Å, and $k = 10$ KNN degree control with sequential edges enabled. We configured EGT with 3 layers having a hidden dimension of 256. During training, we applied

Table S4: Loss weight ratio used in pretraining across experiments.

| Loss term | Symbol | Weight | Notes |
|---|---|---|---|
| Coordinate reconstruction | $\lambda_X$ | 0.1 | Mean square deviation between generated and original coordinates. |
| Intra-block distance | $\lambda_D$ | 0.1 | Inter-atomic intra-block (only applied to residues and nucleotides) distances, and all-by-all small molecular atom-pair distances |
| Atom bond angle | $\lambda_A$ | 0.1 | Over small molecular bond triplets |
| Block/Atom type classification | $\lambda_{\text{EGT\_cls}}$ | 0.01 | EGT-guided block/atom token cross-entropy |
| Masked block/atom classification | $\lambda_{\text{mask}}$ | 0.1 | Recovery of corrupted block/atom tokens |
| Auxiliary label regression | $\lambda_{\text{aux}}$ | 0.1 | Huber loss; $\delta = 1.0$ for Rosetta minimized energy and SASA & HSE features, binding affinity values, and $\delta = 0.2$ for DockQ |

random rotations to the structure coordinates. We kept the pretraining loss weights constant; they are summarized in Supplementary Table S4. All auxiliary regression heads in the training phase shared the same architecture of a two-layer MLP with hidden size 256 and dropout 0.1, and each was instantiated only when its corresponding label type had at least 1,000 training samples.

##### S4.3 Varied Hyperparameters

We selected the hidden dimension of Mamba $d_{\text{model}} \in \{512, 1024\}$, the encoder depth in $\{8, 12\}$, and the decoder depth in $\{4, 6\}$. Quantization configurations varied only in the specific ablation study, with two single-layer FSQ settings ($4^6$ and $5^8$) and four RFSQ settings: two layers with levels $[7, 5, 5, 5]$ or $[8, 5, 5, 3]$, three layers with levels $[5, 5, 3, 3]$, and four layers with levels $[4, 3, 2, 2]$.

##### S4.4 Finetuning Configuration

The pMHC-specific structure generation, classifiers, and optimization settings are provided in Supplementary Section S7. For DockQ and protein–ligand scoring, finetuning froze the pretrained encoder and quantizer and trained only the selected head, using AdamW with a learning rate of $10^{-4}$, a weight decay rate of $10^{-4}$, and a cosine schedule down to $\eta_{\text{min}} = 10^{-6}$. The DockQ head combined a Huber regression term ($\delta = 0.2$) with Pearson and Spearman ranking terms, each at weight 1.0. The MDN head of binding-affinity prediction used 10 Gaussian components, a 256-dimensional hidden layer, and a 10 Å interface-edge cutoff for both training and evaluation. The finetuning objective function is the summation of the density negative log-likelihood (weight 1.0), a Pearson ranking term (weight 0.7), and a listwise ranking term (weight 1.0) against normalized targets $pK$. For StructTokenBench, we adopted the original benchmark procedure and plugged only the frozen OmniScore backbones into the benchmark task-specific procedures.

##### S4.5 Masking Paradigm

We apply the following rule to both atom tokens and block tokens in the independent masking scheme similar to [15]. For a selected token row $i \in \mathcal{M}$, the corrupted token embedding follows the 80/10/10 rule:

$$\tilde{\mathbf{e}}_i = \begin{cases} \mathbf{e}_{\text{mask}}, & \xi_i < 0.8, \\ \mathbf{e}_{\text{rand}}, & 0.8 \le \xi_i < 0.9, \\ \mathbf{e}_i, & 0.9 \le \xi_i \le 1, \end{cases} \qquad \xi_i \sim \text{Uniform}(0, 1). \tag{5}$$

Here, $\mathbf{e}_i$ is the original token embedding, $\mathbf{e}_{\text{mask}}$ is the mask-token embedding, and $\mathbf{e}_{\text{rand}}$ is a randomly selected token embedding.

Optional coordinate masking corrupts the coordinate row while keeping the clean coordinate as the reconstruction target. For a selected coordinate row $i$, we use

$$\tilde{\mathbf{x}}_i = \begin{cases} \bar{\mathbf{x}}, & \xi_i < 0.8, \\ \mathbf{x}_i + \boldsymbol{\epsilon}_i, & 0.8 \le \xi_i < 0.9, \\ \mathbf{x}_i, & 0.9 \le \xi_i \le 1, \end{cases} \tag{6}$$

where $\bar{\mathbf{x}}$ is the centroid over $\Omega_{\text{atom}}$, $\mathbf{I}_3$ is the $3 \times 3$ identity matrix, and $\eta$ controls the relative noise scale:

$$\boldsymbol{\epsilon}_i \sim \mathcal{N}(\mathbf{0}, \sigma_X^2 \mathbf{I}_3), \qquad \sigma_X = \eta \sqrt{\frac{1}{|\Omega_{\text{atom}}|} \sum_{j \in \Omega_{\text{atom}}} \|\mathbf{x}_j - \bar{\mathbf{x}}\|_2^2}. \tag{7}$$

As the target remains the original token, the token recovery objective is:

$$\mathcal{L}_{\text{mask\_cls}} = -\frac{1}{|\mathcal{M}_a|} \sum_{i \in \mathcal{M}_a} \log \hat{p}^a_{i,a_i} - \frac{1}{|\mathcal{M}_b|} \sum_{i \in \mathcal{M}_b} \log \hat{p}^b_{i,b_i}, \tag{8}$$

where $\mathcal{M}_a$ and $\mathcal{M}_b$ are the selected atom-token and block-token row sets, respectively, and $\hat{p}^a_{i,a_i}$ and $\hat{p}^b_{i,b_i}$ are the predicted probabilities of the original tokens.

#### S5 Additional Results

##### S5.1 Effect of ligand-containing structures on protein-only downstream tasks

We next asked whether pretraining on structures that contain small molecules and ligands benefits downstream tasks defined purely on proteins (Fig. S1). Antibody– and nanobody–antigen quality assessment served as the test case, evaluated under Pearson and Spearman correlation, AUROC, and AUPRC. Two pretraining corpora were compared at three model sizes: the full corpus, and an only-protein corpus from which all small molecules and ligands were excluded (Fig. S1). Removing ligand-containing structures did not reduce downstream performance. The highest Pearson correlation was obtained by OmniScore-84M pretrained on only-protein data at 0.916, compared with 0.890 for the same architecture on full data. The 34M pair moved in the same direction (0.895 versus 0.873), and the 24M pair was closer (0.9103 versus 0.901). Spearman correlation and both AUROC and AUPRC values were nearly indistinguishable across the six models, spanning 0.816–0.829, 0.974–0.988, and 0.981–0.989. The single setting that favored full-data pretraining was OmniScore-24M, where AUROC decreased from 0.988 to 0.974 after excluding ligands. Each value comes from a single run without uncertainty estimates, so the small margins should be read cautiously. Within the evaluated settings, adding ligand-containing structures gave no measurable benefit on this task.

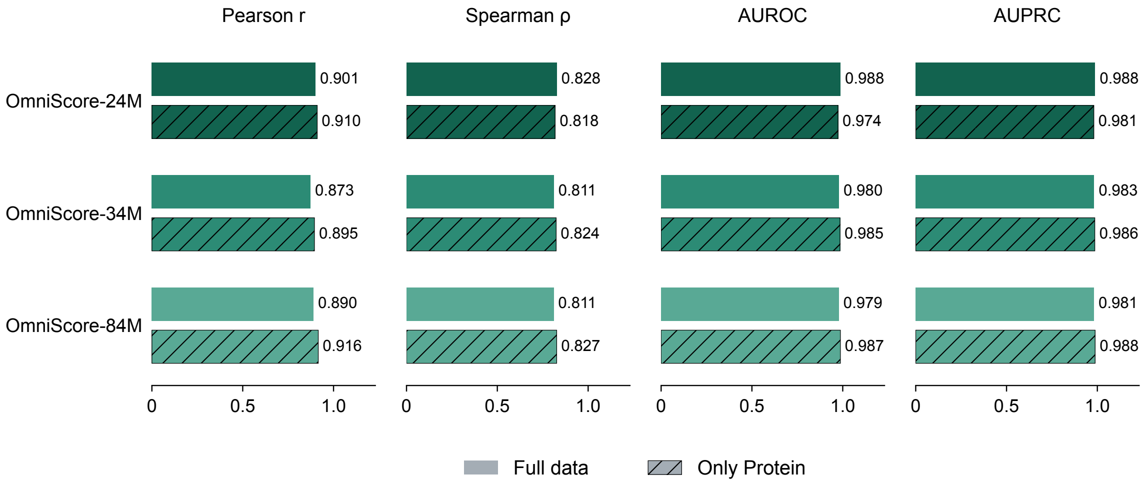


Figure S1: Full-data versus only-protein pretraining for antibody/nanobody–antigen quality assessment. Three OmniScore model sizes (24M, 34M, and 84M parameters) were each pretrained twice: on the full corpus of protein, nucleic-acid, and small-molecule structures ("Full data", solid bars), and on a corpus with all small molecules and ligands removed ("Only Protein", hatched bars). Pretrained models were fine-tuned to predict DockQ, a continuous $[0, 1]$ measure of interface quality, and evaluated on antibody–antigen complexes [7]. Model performances were compared on four metrics: Pearson correlation, Spearman correlation, AUROC and AUPRC.

#### S6 RNA structure ranking results

Table S5: RSR results on RNA Puzzles 14 (bound and free), 18, and 21. OmniScore results use parameter version v3. Each group value is the Spearman correlation between the method score (predicted RMSD for learned regressors) and true RMSD within that candidate group. Mean RS (3 targets) first averages the bound and free correlations for Puzzle 14, then averages the correlations for Puzzles 14, 18, and 21 with equal target weights. Mean RS (4 groups) is the unweighted average across the four candidate groups. Correlations and means are reported to three decimal places; rounding of group values can produce small differences when means are recomputed. Bold values indicate column maxima. The ATOM3D 3DCNN and GNN baselines were retrained on RSR and evaluated as three-seed ensembles [22]; ARES uses released weights [28]. Centered probes remove each training target's feature and RMSD means before ridge fitting (Equation 12); global probes fit without target centering.

| Method | Mean RS (3 targets) | Mean RS (4 groups) | P14 bound | P14 free | P18 | P21 |
|---|---|---|---|---|---|---|
| **OmniScore** | | | | | | |
| Ridge (post-quantization CLS + mean) | −0.191 | −0.193 | −0.507 | 0.107 | −0.210 | −0.163 |
| MLP (post-quantization CLS + mean) | −0.163 | −0.120 | −0.412 | 0.429 | −0.007 | −0.490 |
| MLP (pre-quantization mean) | 0.196 | 0.149 | 0.097 | −0.087 | 0.057 | 0.527 |
| MLP (pre-quantization CLS) | −0.114 | −0.025 | 0.110 | 0.371 | 0.205 | −0.787 |
| Linear probe (pre-quantization mean, global) | 0.094 | 0.164 | 0.377 | 0.367 | 0.112 | −0.200 |
| Linear probe (EGT, centered) | 0.261 | 0.289 | 0.487 | 0.255 | 0.085 | 0.328 |
| Linear probe (EGT, global) | 0.198 | 0.208 | 0.510 | −0.031 | 0.057 | 0.298 |
| Linear probe (Mamba layer 8 mean, centered) | 0.070 | 0.128 | 0.152 | **0.452** | 0.122 | −0.215 |
| GNN (EGT, within-target objective) | **0.423** | **0.427** | 0.526 | 0.350 | **0.337** | 0.492 |
| **Baselines** | | | | | | |
| Rosetta `fa_stack` | 0.169 | 0.157 | 0.255 | −0.009 | 0.158 | 0.224 |
| Rosetta `N_BS` | 0.167 | 0.150 | 0.203 | −0.003 | 0.160 | 0.242 |
| Rosetta `hbond_sc` | 0.125 | 0.112 | 0.075 | 0.073 | 0.071 | 0.229 |
| FARFAR2 Rosetta total score | 0.042 | 0.025 | −0.070 | 0.015 | −0.002 | 0.155 |
| ARES (released weights) | 0.201 | 0.213 | 0.494 | 0.005 | 0.203 | 0.150 |
| ATOM3D 3DCNN (retrained, three-seed ensemble) | 0.274 | 0.267 | 0.363 | 0.134 | −0.031 | **0.605** |
| ATOM3D GNN (retrained, three-seed ensemble) | 0.228 | 0.248 | **0.538** | 0.078 | 0.135 | 0.241 |

**Evaluation set and comparison.** The evaluation removes RNA3DB–RNA-Puzzles overlap and uses evaluation targets absent from encoder pretraining and prediction-head training. Puzzle 14 contributes bound and free candidate groups, while Puzzles 18 and 21 each contribute one group. The three-target mean is the primary summary, so Puzzle 14 does not receive twice the weight of another RNA target. The four-group mean is provided for comparison. With only three RNA-Puzzles targets, differences between methods should be interpreted cautiously. The FARFAR2 baseline uses the stored Rosetta total score; the additional Rosetta rows report individual score terms.

### S7 pMHC interaction classification methods and results

The pMHC sequence pairs, binding labels, and data partitions were obtained from the Qualitative dataset provided in the PREpiBind study [25].

#### S7.1 Structures, backbone, and representation stages

Each complex was predicted with Boltz-2 using multiple sequence alignments (MSAs) for the MHC-II $\alpha$ and $\beta$ chains and no MSA for the peptide, with three recycling steps and 200 diffusion steps. A single structure was predicted per complex, containing three chains: MHC-II $\alpha$, MHC-II $\beta$, and a 15-mer peptide. We used the frozen pretrained OmniScore-24M encoder to embed each complex once; only the downstream classifiers were trained. For every bead-atom, we extracted G (128-dimensional EGT output $\mathbf{H}^G$, before Mamba), P (512-dimensional continuous Mamba output $\mathbf{H}$, before quantization), and E (512-dimensional quantized Mamba latent $\mathbf{Z}$, after quantization), as compared in Supplementary Table S6. Classification (CLS), end-of-sequence (EOS), and padding rows were excluded from all pooling operations.

#### S7.2 Boltz-2 confidence features

The confidence baseline and fusion runs used the per-complex Boltz-2 confidence JSON. Of the 21 raw fields, we excluded five that were uninformative in this setting: `ligand_iptm`, which was identically zero; `protein_iptm`, which duplicated `iptm`; and the three diagonal `pair_chains_iptm_k_k` fields, which duplicated the corresponding `chains_ptm_k` fields. The remaining 16 features were:

- `confidence_score`, `ptm`, `iptm`, `complex_plddt`, `complex_iplddt`, `complex_pde`, and `complex_ipde`;
- the three `chains_ptm_k` fields for $k \in \{0, 1, 2\}$;
- the six off-diagonal `pair_chains_iptm_i_j` fields for $(i, j) \in \{(0, 1), (0, 2), (1, 0), (1, 2), (2, 0), (2, 1)\}$.

#### S7.3 Interface definition

The interface was defined geometrically using an 8 Å cutoff. For the MLP, the interface atom set comprised all MHC bead-atoms within 8 Å of any peptide bead-atom and all peptide bead-atoms within 8 Å of any MHC bead-atom. For the GAT, we used the corresponding interface residue set, comprising contacting peptide residues and MHC groove residues. Approximately 96% of the atoms contributing to whole-complex pooling belonged to MHC, whose structure is nearly identical across complexes sharing an allele. Interface pooling therefore focused the representations on peptide–MHC contacts relevant to binding classification.

#### S7.4 Classifiers

**Run 1: confidence only.** The 16 confidence features were classified using a panel of scikit-learn models: logistic regression, random forest, histogram gradient boosting, and an MLP with hidden-layer sizes $(256, 64)$. Models were selected by validation AUROC; the MLP performed best and provided the reported confidence-only result.

**Run 2: OmniScore + GAT.** Bead-atom embeddings were mean-pooled within each interface residue to form node features. A residue graph connected the $k = 10$ nearest neighbours based on residue centroids. The classifier comprised two GATConv layers with 128-dimensional outputs, four attention heads in the first layer and one in the second, ELU activations, and dropout of 0.2, followed by global mean pooling and a linear logit output.

**Run 3: OmniScore + MLP.** The selected representation (E, P, or G) was mean-pooled over interface atoms to yield one vector per complex. This vector was passed to an MLP with two hidden layers of 256 and 128 units, GELU activations, dropout of 0.2, and a single logit output.

#### S7.5 Confidence fusion

The E+, P+, and G+ variants incorporated the 16 confidence features by feature concatenation. For MLP fusion, the confidence features were concatenated with the interface-pooled OmniScore vector and classified by an MLP. For GAT fusion, the confidence features were concatenated with the graph readout after global mean pooling and passed through an MLP head with a 128-unit hidden layer and a single logit output. The GAT and fusion head were trained jointly from scratch while the OmniScore encoder remained frozen.

##### S7.6 Training and evaluation

We reused the original PREpiBind partitions [25]: a 70:30 training–test split stratified by binding label and MHC serotype, with Fold 0 of the training rows used for validation. Identical peptide/MHC-$\beta$ entries were kept within one partition. This produced 90,328 training, 22,543 validation, and 48,352 test complexes, with approximately 58% positive examples. All features were standardized using means and standard deviations computed from the training set only.

The PyTorch heads for runs 2 and 3 and GAT fusion were trained with AdamW, a learning rate of $1 \times 10^{-3}$, weight decay of $1 \times 10^{-2}$, binary cross-entropy, and a batch size of 128. Training ran for up to 60 epochs with early stopping after five epochs without improvement in validation AUROC; the checkpoint with the highest validation AUROC was restored for test evaluation. The scikit-learn classifiers for run 1 and MLP fusion were likewise selected by validation AUROC. All runs used seed 0 and were evaluated by test AUROC.

Table S6: Test AUROC for pMHC interaction classification comparing NetMHCIIpan-4.3, MixMHC2pred-2.0, Boltz-2 confidence metrics, and OmniScore embeddings on the full test set of the Qualitative dataset from PREpiBind [25]. E denotes post-quantization representations; P denotes pre-quantization Mamba outputs; G denotes EGT outputs before Mamba. A plus sign denotes added Boltz-2 confidence features. The pretrained OmniScore-24M backbone is kept fixed while the task-specific classifiers are trained. GAT uses residue-level graph features followed by global mean pooling; MLP uses atom-level interface mean pooling. Bold denotes the highest AUROC. Uncertainty estimates are not available.

| **Run** | **Whole structure/ interface** | **Feature used for classification** | **Embedding**<br>1. E → post-quantization<br>2. P → pre-quantization<br>3. G → EGT output (before Mamba) | **Test AUROC** |
|---|---|---|---|---|
| NetMHCIIpan-4.3 | — | — | — | 0.838 |
| MixMHC2pred-2.0 | — | — | — | 0.815 |
| 1. Boltz-2 confidence metrics | Whole structure | 16 metrics | — | 0.7801 |
| 2. OmniScore + GAT | Interface | Residue graph; global mean pooling | E | 0.6187 |
| | | | P | 0.8707 |
| | | | G | 0.8698 |
| | | | E+ | 0.7634 |
| | | | P+ | 0.8817 |
| | | | G+ | **0.8821** |
| 3. OmniScore + MLP | Interface | Interface atoms; mean pooling | E | 0.5697 |
| | | | P | 0.7918 |
| | | | G | 0.8266 |
| | | | E+ | 0.7638 |
| | | | P+ | 0.8298 |
| | | | G+ | 0.8487 |
